# Competition between phage-resistance mechanisms determines the outcome of bacterial co-existence

**DOI:** 10.1101/2022.07.11.499539

**Authors:** Olaya Rendueles, Jorge A.M. de Sousa, Eduardo P.C. Rocha

## Abstract

Many bacterial species carry multiple prophages. Despite their potential cost, these elements can provide multiple fitness advantages to the host, such as the elimination of direct competitors. However, the long-term efficiency of prophage induction to displace competitors has received little attention. We experimentally coevolved a polylysogenic *Klebsiella pneumoniae* strain (ST14) with a phage-sensitive strain (BJ1) in several environments resulting in different phage pressures. We then followed the adaptation process and the emergence of resistance. After 30 days, population yield remained stable, and although BJ1 was present in all conditions, its frequency was higher when phage pressure was stronger. Resistance to phages emerged quickly through mutations that prevent capsule biosynthesis. In contrast to our expectation, lysogenic conversion was rare and costly because new BJ1 lysogens exhibited exacerbated death rates and were easily outcompeted. Unexpectedly, the adaptation process changed at longer time scales, where BJ1 populations adapted by fine-tuning the production of capsule, reducing the ability of phage to absorb, while remaining capsulated. These resistant clones are pan-resistant to a large panel of phages. Most intriguingly, some clones exhibited transient non-genetic resistance to phages. Our experimental and modelling results highlight the diversity, dynamics and competition between phage-resistance mechanisms during coevolution and how these are driven by phage pressure.

## INTRODUCTION

Parasites shape the life history and fitness of their hosts. They also impact community structure via predation and competition, and thereby affect numerous ecological and evolutionary processes ^1–3^. Bacteriophages (phages) are very abundant predators of bacteria ^4,5^. Temperate phages either follow a lytic cycle in which they replicate within bacterial cells and release infectious virions, or a lysogenic cycle in which they integrate the bacterial genome and replicate with it. Nearly half of the sequenced bacterial genomes are lysogens ^6^. The dual lifestyle of temperate phages is costly, but can also provide the host with multiple advantages. During lysogeny, prophages may increase biofilm formation ^7^, phosphate acquisition ^8^, or express virulence factors ^9–11^. Inactivated prophages leave genes in the genome that are co-opted by the host and result in functional innovation, e.g. as bacteriocins used in bacterial warfare ^12–14^. Prophages also protect bacteria from closely related phages, a process called superinfection resistance ^15^. Furthermore, when the lytic cycle is initiated in a small subpopulation, it may facilitate colonization by directly mediating competition within communities ^16,17^, because the released virions will infect and lyse closely-related but not identical strains. This can promote the acquisition of adaptive traits from bacterial competitors ^18^. Hence, it is suggested that prophage induction affects bacterial population dynamics, community structure, and evolution ^19–23^.

*Klebsiella pneumoniae* (Kpn) is a ubiquitous enterobacterium, of which, at least 75% of the species’ genomes are polylysogenic, with an average of six prophages per genome ^24^. Many of these are released into the environment and are able to infect other strains ^24^. Phage infection in *K. pneumoniae (Kpn)* is dependent on the presence of the polysaccharidic capsule ^24–27^. Most *Kpn* prophages have narrow host ranges because they only infect strains with a capsule serotype similar to that of their previous host. Nevertheless, the abundance of prophages in the genomes of Kpn and the ability of some of them to infect other strains ^24^, suggests that they play an important role in *Klebsiella* evolution and ecology.

How parasite pressure may alter co-evolving bacterial populations has been seldom addressed, and most of these studies focused on virulent phages ^28–32^. A few other studies have tested the impact of coevolution between lysogens and non-lysogens and the advantages the former provide *in vivo* by mediating bacterial interactions ^17,21,33–35^. However, the relevance of poly-lysogeny for population dynamics during hundreds of generations remains unknown. Studies in *Klebsiella spp.* point out that specific capsule serotype is often required for phage infection ^24,36^, and thus the dynamics of capsule loss influence the phage infection dynamics. Here, we co-evolve two natural isolates of *Klebsiella pneumoniae* both of which have a K2 capsule serotype but are phylogenetically distant; i) the hypervirulent BJ1 strain without inducible or criptic prophages, that was isolated from a liver abscess (ST380) and ii) a polylysogenic multidrug resistant *K. pneumoniae* strain (ST14) isolated from a urinary tract nosocomial infection. ST14 produces multiple infectious virions for which the BJ1 is known to be sensitive ^24^. In this context, cell defence mechanisms, such as restriction modification systems, are not expected to impact population dynamics, as previously shown ^24^. To address if, and how, prophage induction affects the competition outcome between the two strains, we follow their population dynamics through time. We then test for the emergence of phage resistance in the susceptible strain. This reveals the diversity of the emerging mechanisms of phage resistance. It also provides unique insight into how these different mechanisms coexist within a population and evolve through time in response to infection pressure.

## RESULTS

### Temperate phages provide fitness advantage during competition

We first aimed at understanding if the prophages of strain ST14 provide a fitness advantage during competition with phage susceptible strain BJ1. To limit confounding factors such as competition for resources, we grew the cells in a rich environment. To modulate the amount of phage produced, and the ability of the latter to infect, we defined three conditions: (i) LB, (ii) LB supplemented with 0.2% citrate to inhibit phage infection due to calcium chelation ^24,37^ and (iii) LB with mytomicin C (MMC, 0.1 μg/mL) to increase the phage titers in the environment. MMC was added at a concentration that did not significantly affect growth of BJ1 (Figure S1AB), and despite the consumption of citrate by *Klebsiella*, after 24 hours there is still a large amount of citrate remaining, that is sufficient to inhibit infection (Figure S1C). We also quantified the amount of PFU/mL produced by strain ST14 in the different growth conditions. As expected, phage production in ST14 was significantly higher in MMC compared to the two other treatments. Interestingly, ST14 grown in citrate resulted in a marginal increase phage production relative to the control (LB) (Figure S1CD).

To test whether phages could contribute to the competitive fitness of their host, we co-inoculated both strains (BJ1 and ST14) at an initial ratio of 1:1 for 24 hours. First, we tested if mixing the two strains affected the total growth or the population yield. We observed no increased cell death due to the competition in the different growth conditions (Figure S2A). Then, we calculated the competitive index of the strains and observed that there is a large fitness advantage for strain ST14 in all three conditions (Figure 1A). This is most likely because ST14 has both a higher growth rate and population yield than BJ1, except in the presence of MMC (Figure S1AB). Most importantly, we also observed differences in the competitive index depending on the amount of phage released, or its ability to infect the BJ1 (Kruskal-Wallis, dF=2, P=0.006). This effect could be due to the different population yields of each strain in each environment (Figure S2B). To take into account only the fitness effects due to phages, we quantified the strain-interaction effects using C_*i*_(*j*), which measures the effect of mixing two strains *i* and *j* on the viable population size of strain *i*, relative to pure culture controls. This measure accounts for the absolute performance of each competitor in mixed groups (see Methods). Negative *C_i_(j)* values indicate that strain *i* have lower population yield during growth in the presence of *j* than in pure culture, and positive values indicate the opposite. For the phage producer, strain ST14, the competition has no positive or negative effect during competition, most likely because increased release of viruses resulting in ST14 death is outweighed by an exacerbated death of phage-sensitive BJ1 (Figure 1B). In the presence of citrate, a condition in which phage cannot infect, the growth of strain BJ1 is not significantly inhibited. However, in the absence of citrate, when phages can infect, C*i(j)* is significantly lower than zero, indicating a negative effect on the growth of strain BJ1. This effect is dependent on the amount of phage released into the environment, as an increased production of phages by ST14 due to MMC leads to an even lower C_*i*_(*j*) for BJ1 (Figure 1B, Kruskal-Wallis, dF=2, P=0.007).

**Figure 1.**
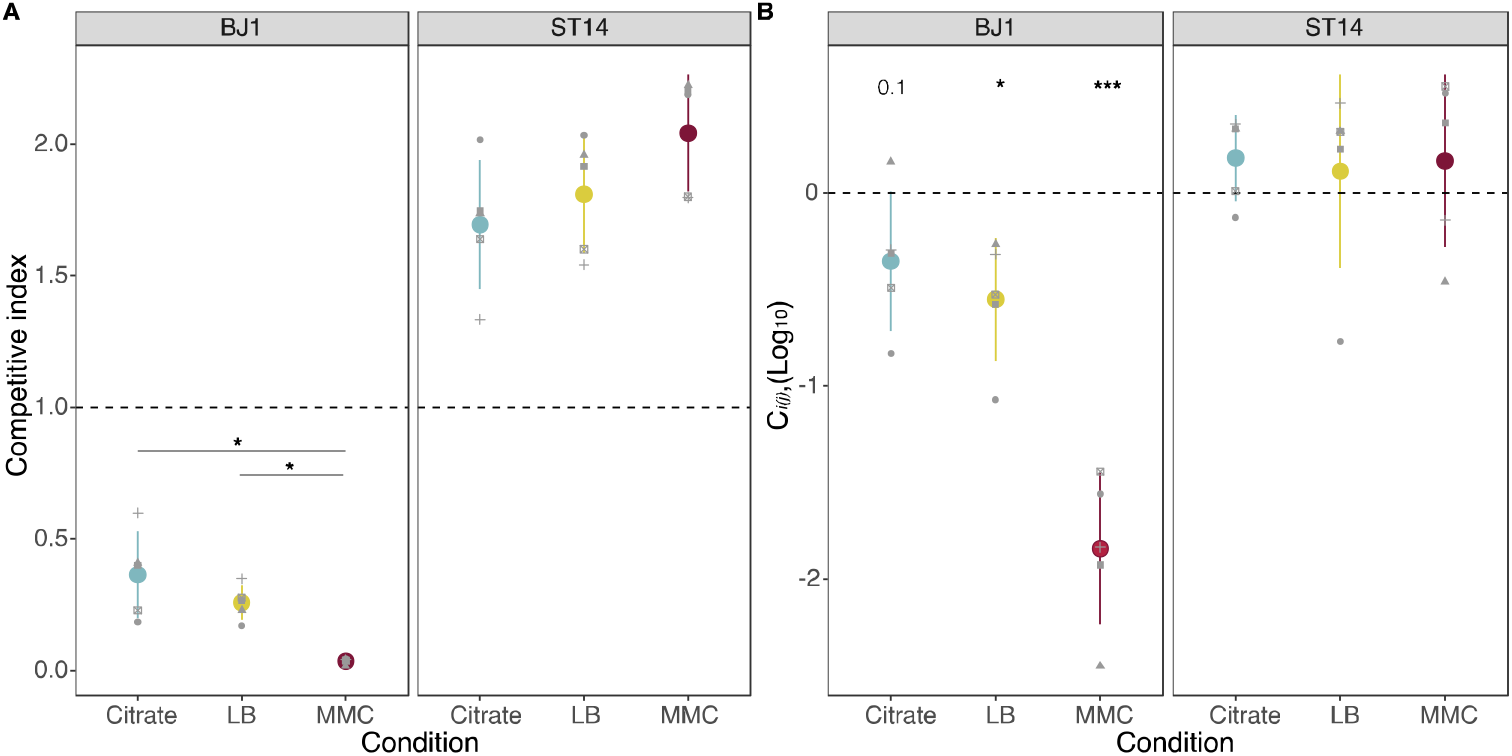
Fitness of strains during competition. **A.** The competitive index is calculated as the final frequency of each strain divided by the initial frequency in the mixed cocultures. * P <0.05, Wilcoxon rank sum test adjusted by Benjamini-Hochberg correction. **B.** The effect of mixing two strains during growth in coculture is given as *Ci(j)*, expressed in log_10_, with *i* representing either strain BJ1 or strain ST14. Positive values represent increased cell numbers during coculture than those expected from the pure cultures. P values corresponds to one-sample t-test for difference of 0. * P< 0.05, ***P<0.001. Each dot shape represents an independent experiment, N=5. Error bars indicate the standard deviation.

We also tested whether ST14 could sense the presence of competition, for instance by quorum sensing mechanisms, and induce prophages and the production viral particles ^38^. Our results show that growth of ST14 with spent supernatant of BJ1 did not result in increased viral release (Figure S1CDE). Taken together, our results show that prophages can increase fitness of their host in co-culture by disfavouring the non-lysogens.

### Resistance to temperate phages emerges rapidly during coevolution

To assess whether ST14 prophages could provide a long-term fitness advantage and outcompete non-lysogens, we set up an experiment in which we allowed three independent mixed populations composed of phage-producing ST14 and phage-susceptible BJ1 strains to coevolve during 30 days, in the three previously defined environments (LB, LB supplemented with 0.2% citrate and LB supplemented with MMC). To follow the evolution of each strain, we plated the populations every day on selective media and counted CFU. As expected, no significant changes in the group yield were observed (Figure S3). This is mostly explained because the dominant strain, the phage producer, does not change its population yield (Figure 2A). In contrast, the frequency of BJ1 decreased rapidly during the first 4 days, suggesting a large initial fitness disadvantage of this strain. This is observed in all three conditions, but it is accelerated in conditions in which phage release is exacerbated (with MMC) and bacterial infection is not restricted (without citrate). Interestingly, through days 4 to 14, evolved BJ1 populations seem to stabilize in number at *ca.* 10^6^ CFU/mL, except for one population evolving in MMC (which increases significantly in frequency). This suggests the emergence of phage resistance. However, after day 15, populations evolving in MMC remain stable whereas the others seem to suffer a second drop in number until day 22 beyond which they once again stabilize at *ca* 10^3^ CFU/mL. Taken together, in 30 days of coevolution, ST14 did not completely displace BJ1 from the populations, even in conditions where the former’s phage induction is exacerbated. This suggests that prophage-mediated competition can be counter-balanced by the evolution of resistance mechanisms in the competitor strain.

**Figure 2.**
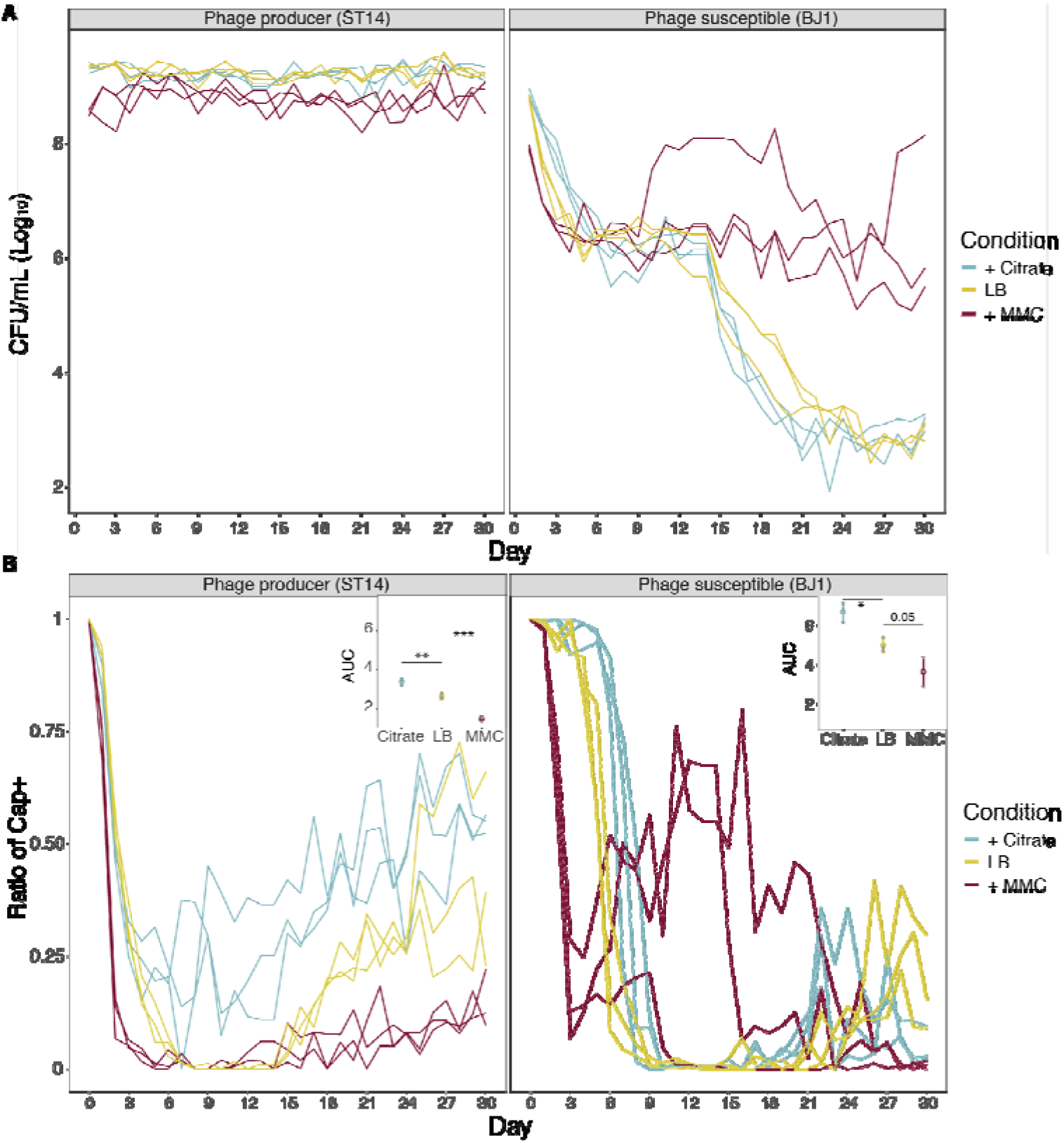
Population yield and proportion of capsulated clones of the two strains during the coevolution experiment. **A.** Total CFU per mL of each strain as estimated every day on selective media. Each line represents an independent coevolving population. **B.** Emergence of non-capsulated mutants in each strain. The insert shows the area under the curve (AUC) during the first 9 days of evolution, as calculated by the function *trapz* from the R package pracma. * P< 0.05,** P< 0.01,***P<0.001 for ANOVA with Tukey *post hoc* corrections.

### Phage pressure drives capsule inactivation as a mechanism of resistance

Our previous results showed that capsule loss provides resistance to phages and faster growth in rich media even in the absence of phages ^39^. Accordingly, throughout the daily plating of coevolving populations, we observed the rapid emergence of non-capsulated clones in all independent populations across the three treatments (Figure 2B). Since capsule inactivation leads to phage resistance ^24–27^, we tested if non-capsulated clones could be under stronger selection for capsule loss when phage pressure is higher (higher density of phages). We observed that the emergence of non-capsulated clones in the BJ1 background is exacerbated in the environment in which phage pressure is greater (insert, Figure 2B), and is diminished when phages cannot infect. Hence, phage pressure accelerates capsule inactivation. Interestingly, this is also the case for ST14, the phage producer, which we had previously shown to be mildly susceptible to its own phages ^24^. Overall, within the first ten days, at least 50% of the population is composed of non-capsulated mutants.

Previous work showed that most of the capsule mutants accumulated mutations in the *wcaJ* gene, the first glycosyltransferase of the capsule biosynthesis pathway ^40,41^. Sequencing of *wcaJ* in the evolved clones (both from BJ1 and ST14 backgrounds) revealed that all but two non-capsulated clones had mutations in *wcaJ*, most of which resulted in a loss-of-function (Table S2). Specifically, 16 out of the 18 non-capsulated ST14 clones analysed had the same mutation, namely a thymine (T) insertion at position 540, in a stretch of 4 T residues, which resulted in a frameshift. However, *wcaJ* mutations in the 36 evolved clones of evolved BJ1 were more diverse, with several different mutations even across individual clones isolated from the same population. Mutations in fourteen clones resulted in either a frameshift or a premature stop. Two different mutations affected the same aminoacid, a tyrosine in position 339. Taken together, as previously shown, capsule inactivation mostly emerges by mutations in *wcaJ.*

### The emergence of new lysogens is rare and potentially unstable

Some resistant clones of strain BJ1 are capsulated, which lead us to hypothesize that they evolved other resistance mechanisms. To test this, we analysed at different time points the resistance mechanisms of the capsulated clones in the population. We expected to find BJ1 lysogens, since super-infection exclusion due to the lysogenization of capsulated bacteria could prevent further infection by the same phages. Further, our previous work had already shown that, when infected with phage lysate at high titers, at least two of the four intact phages from strain ST14 could lysogenize BJ1 ^24^. To quantify the proportion of lysogenized BJ1 cells, relative to other resistance mechanisms, we isolated over 1200 capsulated clones at different time points (Figure S4). We identified the clones that were resistant to purified phage lysates of strain ST14 and that produced phages when exposed to MMC in our culture conditions. More precisely, we analysed the differences in the area under the growth curve of each clone, both when they were grown in LB (control), when phage lysate was added (to distinguish between resistant or susceptible), and when MMC was added (to induce prophages and identify newly lysogenic clones). Together with the resistant non-capsulated clones (Figure 2B), this provides a detailed overview of the different mechanisms of resistance, their proportion, and their temporal dynamics throughout the experiment (Figure 3).

**Figure 3.**
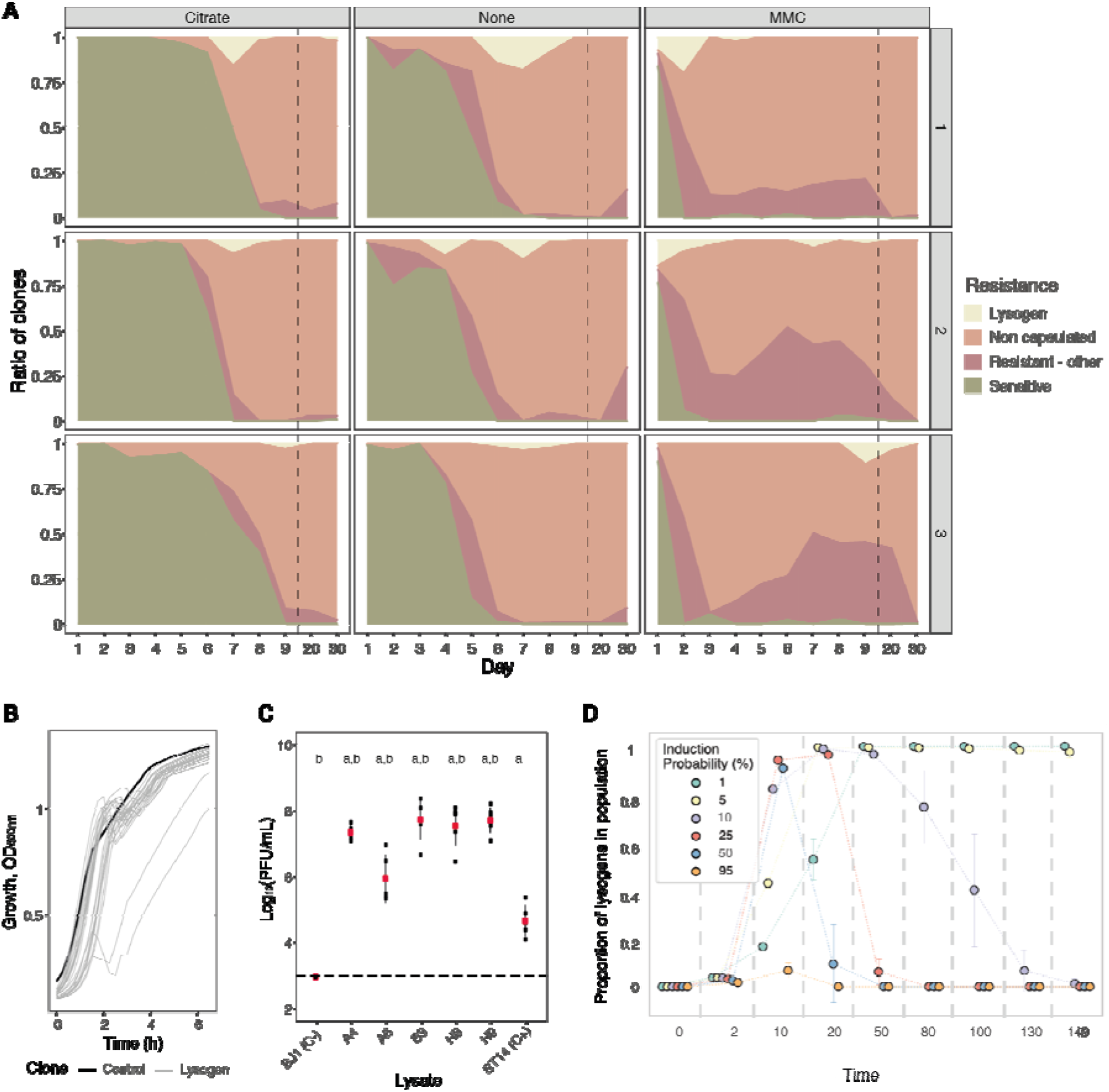
Evolution of resistance mechanisms in strain BJ1. **A.** Ratio of clones from each coevolving population that are susceptible (green), non-capsulated (light pink), capsulated lysogens (beige), or capsulated but resistant by other undefined mechanisms (dark pink). N.B. Dashed line indicates when x-axis, no longer follows a linear scale. **B.** Growth of newly lysogenized clones reveal significant death during exponential phase (in the absence of induction), as measured by the optical density. Black line corresponds to the control, BJ1 ancestor. All independent growth curves, with and without induction are represented in Figure S5. **C.** PFU/mL produced without induction by five selected new lysogens derived from BJ1 and isolated at day 1 for A4 and A6, at day 4 for B3 and day 9 for H8 and H9. Dashed line indicates the limit of detection of the essay. Each black dot represents an independent strain lysate and large red dots represent the mean. Error bars correspond to the standard deviation. Two-sided t-test ‘a’, P<0.001 compared to ancestor BJ1 (negative control, C-) and ‘b’, P<0.05 compared to ST14 (phage producer, positive control, C+). **D.** Simulated temporal dynamics of proportion of lysogens in the populations, as calculated by eVIVALDI. Each circle corresponds to the central tendency of replicate simulations, with the different colours indicating a given probability of spontaneous prophage induction (shown in the legend, values approximated to nearest major integer). The error bars correspond to the standard deviation across the replicate simulations. In the represented simulations, the probability of acquisition of a phage resistance mutation (capsule loss) is 0.001, and the fitness cost of this mutation is 10% of the bacterial growth rate, as calculated in ^39^.

We first observed that the proportion of susceptible clones quickly decreases and is dependent on the amount of phage able to infect in the population (Figure 3), susceptible cells being scarcer in populations evolved on MMC. Specifically, most tested clones were resistant by day 2, 6 and 8 in populations evolving in MMC, LB and citrate, respectively. As expected, lysogens emerged in all populations, but remain in low proportion and their numbers quickly dwindle after their emergence (Figure 3). We verified that the 94 identified lysogens, out of the 1209 clones we screened, were *bona fide* lysogens as shown by their production of phages that infect naïve BJ1 cells, both when induced by MMC (92 out of 94), and in the absence of induction (87 clones out of 94) (Figure S5).

Interestingly, we observed that when new lysogens are grown in LB, in the absence of induction, there is a detectable amount of cell death, and growth delay at the end of exponential phase in at least in 29 out of 94 tested clones (Figure 3B and S6). This could correspond to a high frequency of spontaneous induction in the newly lysogenized bacteria. Indeed, we observe a large amount of phage release, as evidenced by large inhibition halos on an overlay of ancestral BJ1. We selected five lysogens that descended from BJ1 and consistently showed large inhibition halos. We quantified the amount of phage released, in the absence of induction, and quantified infection on a lawn of ancestral BJ1. New lysogens produced between 100 and 1000 more PFU/mL than the ancestral phage producer (ST14) (Figure 3C). This suggests that protection by lysogeny results in significant fitness costs (Figure S5 and S6A).

To study the impact of prophage acquisition in the long-term stability of lysogens in a population, we used eVIVALDI, an individual-based model for microbial interactions and evolution ^42^. We used these simulations to explore different rates of induction of prophages, in the presence or absence of abiotic agents. We designed a scenario where a population of initially sensitive bacterial cells is exposed to an inoculum of temperate phages, and we follow the populations for a period of 150 iterations (e.g., approximately 150 generations). Simulated bacteria can either be infected by phage (thus either dying upon a lytic infection or becoming lysogens if the phage integrates the bacterial genome) or become resistant to phage by mutation (*i.e.*, capsule inactivation, which decreases their growth rate). We then measured, over time, both the total number of cells and the proportions of lysogens.

We observed two main patterns. When prophages have low spontaneous induction rates (1 to 5%), they generate stable, non-costly lysogens. As a consequence, phages spread slowly in the population and this allows time for the phage resistant mutants to emerge and increase to high frequencies. However, because these mutations are costly, lysogens slowly but eventually displace them. This results in a sigmoidal-like temporal frequency of lysogens, where at the end of the simulations most of the resistant population is composed of lysogens (Fig 3D and lower left part of panel S7A). These dynamics are in contrast with the bell-shaped dynamics observed for high or intermediate rates of spontaneous prophage induction (i.e., >=11%), where lysogens quickly invade the population but are absent at the end. Such high rates correspond to unstable lysogens that quickly die due to spontaneous induction of their prophages. These conditions facilitate the propagation of phage throughout the population (due to fast phage amplification), and thus also the result in the rapid emergence of new bacterial lysogens (t=10 in Fig 3D and Fig S7A). If lysogens are protected from new phage infections, becoming a lysogen is an extremely advantageous strategy, but with a very short-term effect: since high rates of induction are very costly for the cell, in the long term lysogeny is counter-selected if non-lysogens can become capsule-less mutants (Fig S7B). As a result, when spontaneous induction rates are high and these mutants emerge frequently and incur in little fitness cost, there will be few, or no lysogens in the populations, as they are expected to be outcompeted (top-right areas for the heatmaps in Fig S7A). This is consistent with our experimental results, where BJ1 clones quickly become lysogens with high induction rates which leads to their removal from the population by the end of the experimental evolution.

In our simulations, the absence of capsule-inactivating mutations (resistance probability = 0, rightmost column of the heatmaps in Fig S7A), implies that populations either become extinct (if induction rates are too high) or are completely composed of lysogens. In contrast, our *in vitro* experiments revealed some resistant clones that were still capsulated and non-lysogens, indicating alternative mechanisms of resistance to phage. These novel clones were more frequent in populations under high phage induction pressure (MMC), when the cost of lysogeny is high, and less frequent under growth in LB with or without citrate (Kruskal-Wallis, dF=2, P=0.03) (Figure 3). Taken together, our results show that most clones became resistant by capsule inactivation, a few by lysogenization, and others by novel mechanisms. They also suggest a strong competition between multiple phage resistance mechanisms.

### Several changes in the capsule production play a role in resistance to phages

To identify the mechanisms of resistance to phages that involved neither capsule loss nor lysogeny, we characterized twelve random clones out of the 328 clones with such profiles. We measured their capsule production and resistance to purified phage lysate, either on a layer of melted agar or during growth in liquid culture. We then tested the ability of each clone to adsorb phage lysate to understand if resistance occurs prior to entering the cell. As controls, we used the ancestral strain (BJ1), as well as an *ΔrcsB* mutant, with reduced capsule expression, and a non-capsulated *ΔwcaJ* mutant (Figure 4). Additionally, we performed whole genome sequencing on all twelve resistant clones and looked for mutational targets, using the ancestral sequence as reference (Table 1).

**Figure 4.**
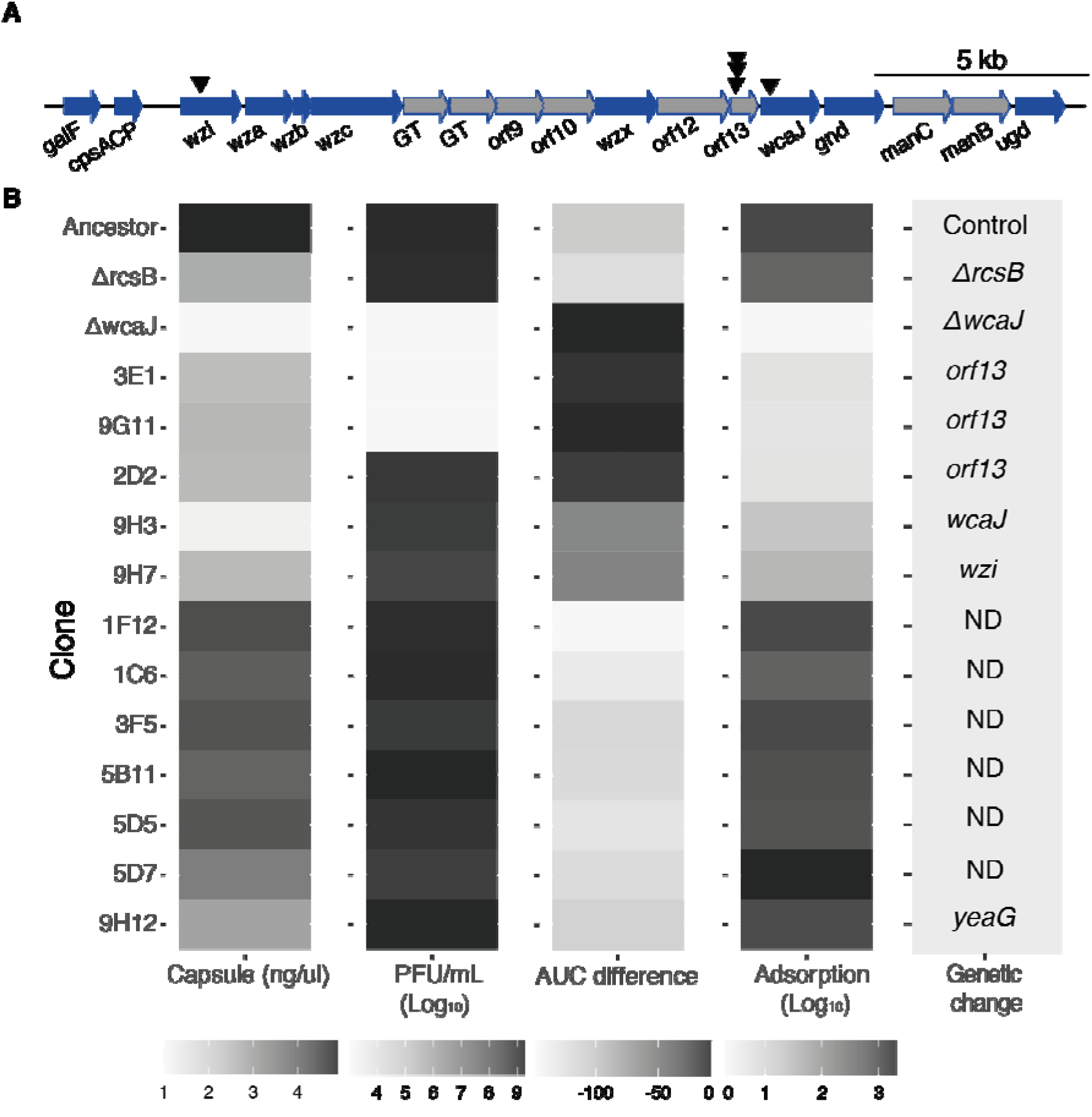
Characteristics of phage resistant clones. **A.** Schematic organization of the capsule operon of strain BJ1. Black triangles represent mutations observed in the capsule operon. Blue arrows indicate core genes common to all *K. pneumoniae* capsule serotypes. Grey arrows correspond to serotype-specific genes. GT stands for glycosyltransferase. The diagram was generated with genoplotR package. **B.** For each clone, we evaluated the amount of capsule produced, the sensitivity to phage on overlay (PFU/mL) and on liquid culture (AUC difference), and the ability of the phage to be adsorbed. The average of three independent replicates is shown. The experiments were performed with three independently generated lysates, when applicable. The AUC difference represents the effect of adding phage to a growth curve. If no effect is observed, the difference in AUC is ~ 0. When phage is added to susceptible clones, the AUC is lower and thus different from the control curve. ND: none detected.

**Table1.** List of mutations identified in the resistant clones sequenced. Location indicates if the mutation is found on the chromosome (C) or plasmid (P). Pop stands for population. The number of mapped sequences is also reported. Clones with less than 98% of mapped sequences are displayed in italics.

| Clone | Pop | Condition | Day | Location | Position | Change | Mutation type | Annotation | Function | Mapped sequences |
| --- | --- | --- | --- | --- | --- | --- | --- | --- | --- | --- |
| 1C6 | 1 | MMC | 1 |  |  |  |  |  |  | 98.3 |
| 1F12 | 2 | MMC | 1 |  |  |  |  |  |  | 98.4 |
| 2D2 | 2 | MMC | 2 | C | 3,994,990 | T→A | I110F (ATT→TTT) | *orf13* | K2 capsule gene; Polysialic acid O-acetyltransferase | 98.2 |
| 3E1 | 1 | MMC | 3 | C | 3,995,203 | (T)<sub>8→7</sub> | coding (115/669 nt) | *orf13* | K2 capsule gene; Polysialic acid O-acetyltransferase | 98.4 |
| 3F5 | 3 | MMC | 3 | C |  |  |  |  |  | 98.6 |
| 5B11 | 1 | LB | 5 | P | 34,553 | A→G | intergenic (+401/-389) |  | hypothetical protein/hypothetical protein | 98.2 |
|  |  |  |  | P | 34,561 | A→G | intergenic (+409/-381) |  | hypothetical protein/hypothetical protein |  |
| 5D5 | 2 | LB | 5 |  |  |  |  |  |  | 98.5 |
| 5D7 | 2 | LB | 5 |  |  |  |  |  |  | 98.6 |
| 9G11 | 1 | LB | 9 | C | 3,995,203 | (T)<sub>8→9</sub> | coding (115/669 nt) | *orf13* | K2 capsule gene; Polysialic acid O-acetyltransferase | 98.3 |
| 9H7 | 2 | Citrate | 9 | C | 2,890,708 | C→T | R121R (CGG→CGA) | *dcuR* | Transcriptional regulatory protein DcuR | 98.5 |
|  |  |  |  | C | 4,009,965 | Δ11 bp | coding (245-255/630 nt) | *wzi* | K2 regulatory capsule gene |  |
| 9H3 | 1 | Citrate | 9 | C | 3,994,389 | C→T | M61I (ATG→ATA) | *wcaJ* | UDP-glucose:undecaprenyl-phosphate glucose-1-phosphate transferase | 98.5 |
| 9H12 | 3 | Citrate | 9 | C | 2,575,329 | C→T | D615N (GAC→AAC) | *yeaG* | Protein kinase YeaG | 98.6 |
|  |  |  |  | P | 23,432 | G→A | R34Q (CGG→CAG) | *soj* | Phage cox protein (PF10743), annotated as plasmid-partitioning |  |

The integration of these analyses revealed several resistance genotypes. Five independent clones had mutations in the capsule operon. Two clones (3E1 and 9G11) had frameshift mutations in a gene coding for an acyltransferase, *orf13*, and potentially leading to a change in the capsule’s biochemical composition (Figure 4A, Table 1). These clones were fully resistant to phage both in liquid and on agar, and had a diminished capsule production, comparable to the phage susceptible *ΔrcsB* mutant. However, phage particles could not successfully adsorb to the surface (Figure 4B). The three remaining clones had a non-synonymous mutation in *orf13* (2D2) and in *wcaJ* (9H3) and an 11 base-pair deletion in the capsule regulator *wzi* (9H7). These clones have reduced capsule expression comparable to mutations 3E1 and 9G11 in *orf13*, reduced phage adsorption and an increased resistance to phage in liquid media (Figure 4B). Surprisingly, these three clones are susceptible to phage when growing on agar. These results suggest that the effect of small capsule modifications in phage resistance might be dependent on the environment. Finally, we found no mutations in known phage defence mechanisms, such as CRISPR-Cas or restriction-modification enzymes. Taken together, our results show that there are multiple paths to resistance that involve modulating either the capsule amount or its composition.

Interestingly, the remaining seven clones that were identified in our initial screens to be resistant seem to be susceptible to phage lysate in all subsequent tests. Despite their marginally lower capsule production, we could not detect mutations in their genome relative to their ancestors (except for one clone with an intergenic mutation). To discard the possibility that this could be due to a problem in our initial screen for resistant clones, we returned to the original glycerol stocks and retested these clones for their resistance to phage during growth in liquid (*i.e.* the same conditions as the screen) (Figure 5). To avoid a possible loss of the phenotype due to culture passaging, we initiated the growth curves directly from the glycerol stock without performing a preconditioning culture, that is, an acclimation step. When the culture reached OD ~0.2, we added the phage lysate. We observed that the cultures grown directly from the stocks were resistant to phage. The difference between the clones from the glycerol stock and those sequenced is that the sequenced clones underwent two extra round of LB passaging without phage pressure. Hence, these results suggest that transient resistance to phages can emerge without mutations (Figure 5 and Figure S8).

**Figure 5.**
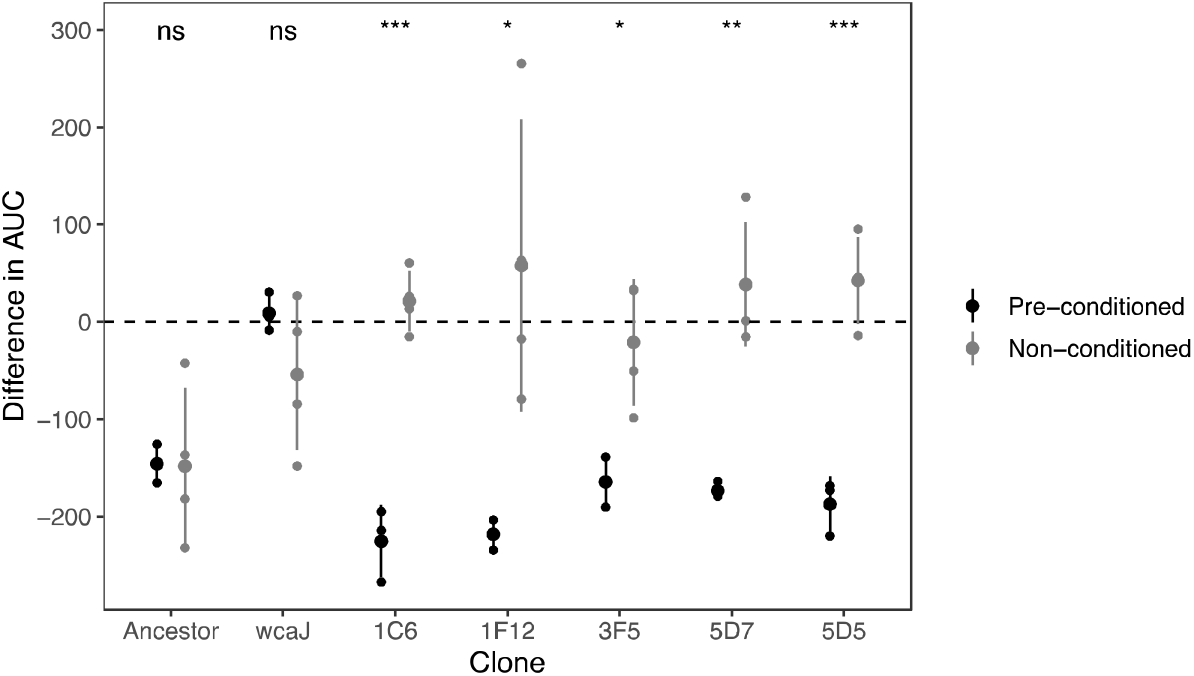
Transient resistance to phages. The difference in the area under the curve (AUC) represents the effect of adding phage to a growing culture as estimated by the difference in growth curve in the absence of phage and with phage (Figure S8). Values below 0 indicate strains are sensitive to phage, and values close to 0 indicate that there is no effect of adding phage to the culture. Non-conditioned clones are those directly grown from glycerol stock, whereas pre-conditioned clones, had been reisolated twice, and grown overnight prior to performing the growth analyses. The ancestor, BJ1, sensitive, and the non-capsulated mutant (*ΔwcaJ)*, are included as controls for the difference in culture conditions. Statistics represent t-tests to check from differences between clones directly from stock and those sequenced (after two passages in LB). ns means non-significative, * P < 0.05, ** P<0.01 and ***P <0.001.

### Capsule modifications but not lysogenization provide cross-resistance to other phages

We sought to test whether resistance to phages from ST14 could result in resistance to phages produced by other strains. To test potential cross-resistance between lysates, we produced phage lysates from strains 03-9138, ST17 and T69, all of which share the same capsule serotype as strain BJ1 (K2) and for which we have previously shown that they successfully infected BJ1 ^24^. We first infected the non-lysogenized BJ1 clones that are capsulated and resistant to ST14 phages. We observed that these clones were also resistant to the phage in lysates of strain 03-9138 and ST17 but not those of T69, suggesting that one of the phages of T69 may not use the capsule as a primary receptor (Figure 6A and Figure S9A). Crossresistance could result from phages sharing a specificity for the capsule serotype ^36^. The strains from which we produced the lysate have very dissimilar prophages, (wGRR<0.25, a measure of phage similarity for all intact phages Figure S9B, ^24^). Yet, 23 proteins from these phages showed sequence identity higher than 50% with proteins present in the two ST14 phages (Figure S9C). The functional analyses of these proteins using pVOG revealed that some are structural proteins potentially involved in infection (*e.g.*, tail proteins) (Figure S9D). They may provide different phages with similar tropism, thereby explaining the observed cross-resistance among lysogens of the same serotype.

**Figure 6.**
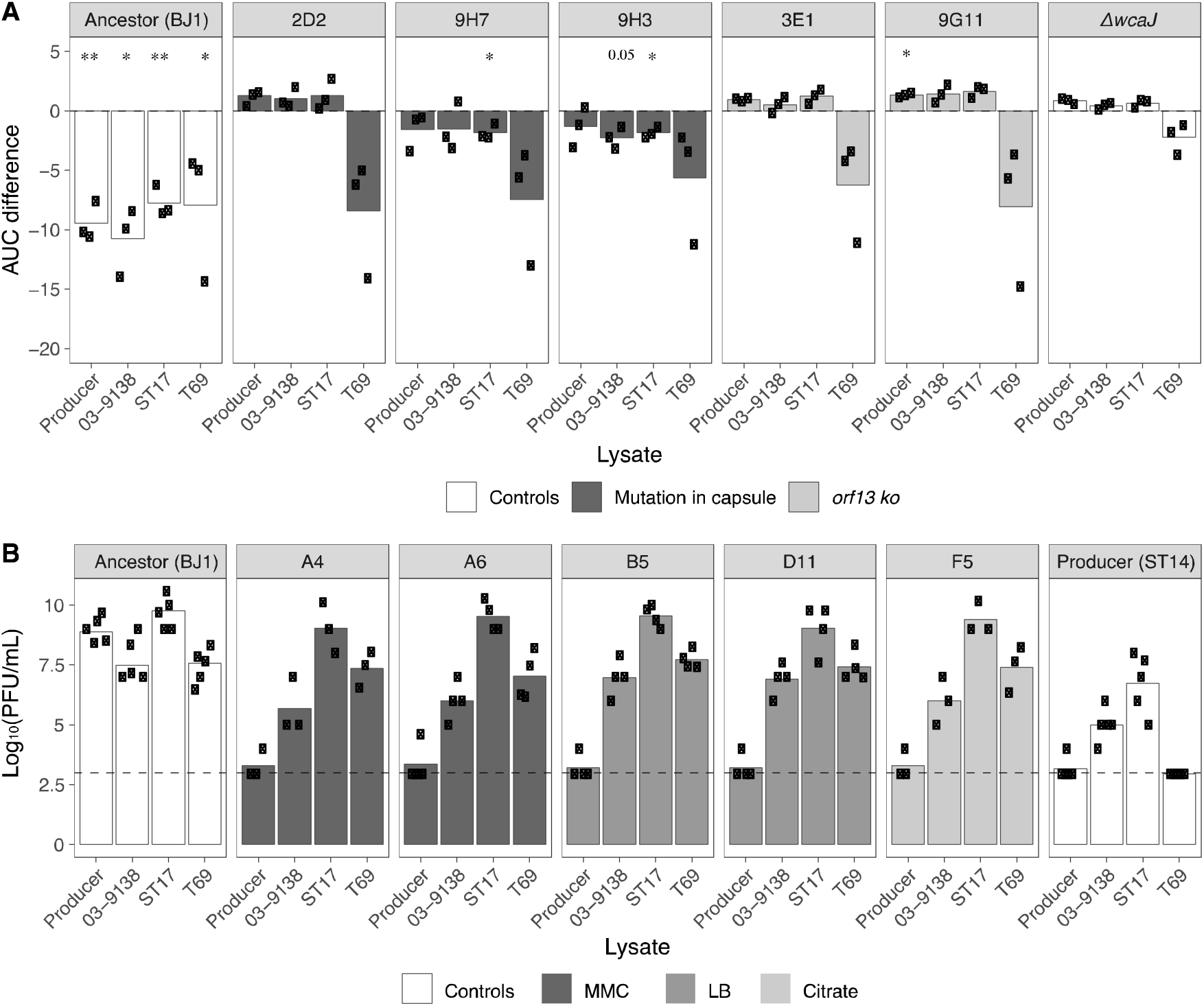
Cross resistance to phages from other lysates. **A.** The area under the curve for capsulated (and non-lysogenized) resistant clones was calculated. The AUC of control cultures with LB was subtracted from those that were challenged with phage lysates. (Growth curves are shown in Figure S9A). Each dot represents an independent assay. One-sample t-tests were performed to test the difference from 0 (growth in LB). * P<0.05; ** P<0.01 **B**. PFU/mL of three independently generated lysates on lawns of new BJ1 lysogens from different evolutionary treatments. Lysogens A4 and A6 evolved in MMC and were isolated at day 1, lysogens B5 and D111 evolved in LB and were isolated at day 6 and 7 respectively, and clone F5 evolved in citrate and was isolated at day 7. Each black dot represents an independent assay. The dashed line represents the limit of detection of the assay.

We then tested whether the lysogenized BJ1 clones, resistant to ST14 lysate, were also resistant to other lysates. To do so, we assessed plaque formation on lysogen lawns rather than growth inhibition, as these new lysogens already exhibited significant cell death due to phage outburst (Figure 3B). Despite similarities between phages across lysates, BJ1 lysogens were only resistant to ST14 lysate, remaining sensitive to lysates produced by other strains (Figure 6B). Taken together, opposite to resistant clones with capsule modifications, integration of ST14 phages into BJ1 does not result in resistance to super infection against a larger array of serotype-specific phages.

## DISCUSSION

We explored how temperate phages drive population dynamics during competition between two *K. pneumoniae* strains. We followed the adaptation of bacterial populations through time under different degrees of parasite pressure by modulating phage infection in two ways: either modifying the density of phages (change in induction rates with MMC) or their ability to adsorb to the bacteria (use of citrate). We hypothesized that phage sensitive BJ1 would coexist for longer time with polylysogenic ST14 in conditions with low phage infection but would be outcompeted faster in conditions of high phage concentration. This was expected based on theoretical works ^43^ and was indeed observed in the initial 24 hours of the competitions, where BJ1 populations decreased in frequency faster under higher phage pressure. In populations with less pressure, both ST14 and BJ1 are present in the population. The co-existence established between the two strains is very dynamic and suggests continuing co-evolution, since we observe an initial drop in phage-susceptible BJ1 populations, a stabilization of its frequency between day 6 and 15, and a second drop in frequency. Under high phage infection regime (MMC environment), BJ1 populations dropped faster than in the other conditions, but after 6 days of co-evolution, the frequencies of both strains reached an equilibrium. This also resulted in a higher frequency of BJ1 strain at the end of the evolution experiment. Such high frequencies in these populations could be due to the emergence of more diverse resistance mechanisms or to lower fitness costs of such resistance. Indeed, capsulated non-lysogenized BJ1 resistant clones emerged more frequently in populations evolving under higher phage infection pressure.

We expected that co-existence between strains under high infection pressure could be the result of extensive BJ1 lysogenization by ST14 phages. Lysogenization often renders bacteria resistant to phages ^44,45^, and has been proposed to limit the period of time where prophages can work as weapons against sensitive bacteria ^43^. Further, we observed in a previous study the frequent lysogenization of BJ1 when exposed to highly concentrated lysate of ST14 ^24^. Surprisingly, here we rarely observe lysogens (Figure 3A). A key difference between these studies is that phage titers are lower here than in the concentrated and purified phage lysates used previously. Experimental ^46^ and modelling studies ^42^ have shown that lysogeny is favoured under high local concentrations of phage, which could explain our results. Another difference among studies is the number of generations, which is much higher in the present study. Here, we do observe the emergence of new lysogens, but this is followed by their disappearance. Both our experimental and modelling results suggest that these new lysogens are less fit probably due to by high rates of spontaneous induction and are thus counterselected to benefit other resistant clones. Our results are also in line with studies with *E. coli* in a murine model, in which high induction rates of lambda phage resulted in decreased fitness of the lysogen ^21^. The presence of a polylysogen could further complicate the acquisition of resistance, because it requires multiple lysogenization events, one for each phage, for full resistance. For instance, it was observed that during a 24-hour competition *in vivo* between a non-lysogenic and a single lysogen strain of *P. aeruginosa*, almost all non-lysogens underwent lysogenic conversion. However, when the same experiment involved a non-lysogenic and a polylysogenic strain (with two phages), the authors observed that less than half of the clones became lysogens ^33^. These and our results suggest that lysogeny is less frequently a mechanism of phage resistance when the rates of spontaneous induction are high or when bacteria are targeted by multiple different phages. In these contexts, evolution can be constrained by competition between the multiple phages. Additionally, it can also be driven by the complex social interactions between newly lysogenized bacteria and bacteria which are resistant by alternative mechanisms.

Here, we show that during the early stages of co-evolution with a close-related strain releasing temperate phages, phage resistance often results from the inactivation of the capsule. This is also observed when *Klebsiella spp* are treated with virulent phages ^25,26,47,48^. Yet, we observed a resurgence of the capsulated clones after several days. This resurgence was also observed in other long-term evolution experiments with BJ1, albeit not in the exact same experimental conditions ^49^. In the latter study, clones of BJ1 evolving in LB also tended to diminish capsule production with time, suggesting that capsule expression can be modulated to reduce production costs. This suggests that complete capsule inactivation may be outcompeted by clones with other mechanisms of resistance that are still able to (at least partially) form a capsule. In agreement with this hypothesis, we observed that some capsulated resistant mutants code for mutations in the capsule operon that result in lower capsule production, which could also limit phage adsorption. Small capsule changes could thus allow escape from phage predation, as most *Klebsiella* phages are serotype-specific^24,36,41^.

We also observed that some phage-resistance mechanisms are dependent on environmental structure. Some mutants were only resistant to phages in liquid media, *i.e.*, in the environment where they evolved. Yet they were fully sensitive to the same phages when growing on agar, which is a highly structured environment. These results are unexpected given previous theoretical work suggesting that spatial structure favours bacterial survival under phage pressure ^50–52^. Differences in resistance across environments are not necessarily linked to a mere reduction in the production of capsule because the *ΔrcsB* mutant that produces less capsule remains sensitive to phage in both liquid and agar (Figure 4). A plausible explanation could be that in well-shaken liquid, phage-bacterium interactions are more sporadic and unstable, compared to the increased stability of interactions in agar. In the latter, or other structured environments, phages have more time, and possibly more opportunity, to stabilize their interaction with bacteria, which could explain why capsule-altered mutants are not fully resistant in these conditions. Finally, allowing coevolution to run for more generations revealed interesting adaptation mechanisms. We show that selection ultimately favours mutations that provide a larger benefit in the evolutionary context rather than others like capsule inactivation which, *in fine*, may be more costly as it remains an important cell surface structure in *Klebsiella.*

Surprisingly, our study revealed that some originally phage-resistant clones reverted to susceptibility after two passages in the absence of phage pressure. Yet, the sequence of these clones failed to reveal any mutations that could explain the changes. Transient resistance has been receiving increased attention, with several recent reports of such phenomena suggesting that this could be a frequent mechanism of resistance. For instance, Hesse and colleagues sequenced 57 different clones of *K. pneumoniae* resistant to a virulent phage and found that almost half of them lacked identifiable mutations ^25^. A plausible genetic basis for these and our resistant clones could be linked to the nucleotide sequence of *wcaJ* of the K2 capsule. These clones may have initially accumulated mutations in simple sequence repeats (SSR) that genome assemblers and variant calling software have difficulties in dealing with. Furthermore, SSR are known mutational hotspots in rapidly evolving traits and their changes are easily reversible ^53^. Indeed, capsule inactivation in ST14 is generally associated with an insertion of a thymine among a repeat of thymine residues (Table S2). However, other, non-genetic mechanisms of transient resistant have also been described. One is the epigenetic-based resistance based on DNA modifications, such as methylation. This was previously shown to regulate the length of the O-antigen length by phase variation in *Salmonella enterica*, and resulted in transient phage resistance ^54^. Additionally, a recent study has shown that cell wall shredding is a transient phenotype that leads to phage resistance in filamentous actinobacteria, *B. subtilis* and *E. coli* ^55^. An analogous process in *K. pneumoniae* could involve the shredding of the capsule or heterogeneity in its production in response to phage pressure, leading to a phenomenon which could be similar to phenotypic resistance in that it is not based in genetic modifications ^56^. This phenotypic resistance is manifested as complete resistance (as there is no initial drop in cells when the phage is added to unconditioned cultures) and could be based on a reduced state of phage adsorption due to changes in capsule. Ultimately, all these transient resistance mechanisms seem to affect the integrity or length of cellular surface structures, be it the LPS, the cell wall or, potentially, other specific cell surface receptor. These non-genetic mechanisms may allow the bacteria to resist phage infection without having to endure a corresponding mutational load caused by the genetic mutations.

Finally, our data is in agreement with a previous study in which the amount of phage present was shown to drive the nature of resistance mechanisms ^57^. At low phage infection pressure, resistance emerges mainly by receptor loss: the pili in *P. aeruginosa* and the capsule in *K. pneumoniae*. Such mutations are costly, as they are constitutive. However, at high phage pressure, inducible resistance like CRISPR-Cas or transient resistance is more advantageous, as the fitness cost is only paid when phage is present ^57^. Taken together, our results highlight the complexity of bacterial interactions, which are shaped by the prophages that reside within cells, and how these may alter evolutionary outcomes by driving the emergence and maintenance of diverse resistance mechanisms.

## MATERIALS AND METHODS

### Bacterial strains and growth conditions

Bacteria were grown at 37° in Luria-Bertani (LB) agar plates or in 4 mL of liquid broth under vigorous shaking (250 rpm). Chloramphenicol (30 μg/ml) and trimethoprim (100 μg/ml) were used to select for strain BJ1 and ST14 respectively.

### Competition calculations

Calculations of *B_ij_, C_i_(j)* were performed as reported in ^58^. *(i) Unidirectional mixing-effect parameter C_i_(j).* The effect of mixing two strains *i* and *j* on the population yield during growth of focal strain *i* was quantified by the one-way mixing effect parameter *C_i_(j).* To calculate this parameter, the expected log10-transformed yield of strain *i* based on pure-culture performance (corrected for the frequency at which strain *i* was added) was subtracted from its actual log-transformed yield during competition with strain *j*.

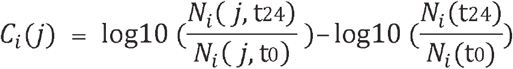

Positive *C_i_(j)* values indicate that strain *i* grew to a higher population size in the experiments in the presence of strain *j* than in pure culture, whereas a negative value indicates that mixing with *j* negatively affected growth yield of *i*. *(ii) Bidirectional mixing effect parameter B_ij_. B_ij_* is the difference between the actual total (log_10_-transformed) group cell count in a mix of strains *i* and *j* and the value expected from pure culture performance of the same strains (corrected for initial frequencies of each strain).

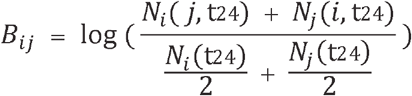

Positive and negative *Bij* values indicate that total productivity is higher or lower, respectively, than expected from pure culture performance.

### Coevolution experiment

Three clones from each strain were used to inoculate overnight cultures, which were then diluted at 1:100 and used to initiate the three independent mixed populations in a ratio 1:1, in a final volume of 4 mL. Each of the three mixed populations evolved in three different environments: (i) LB, (ii) LB supplemented with 0.2% citrate and (iii) LB with mytomycin C (MMC, 0.1 μg/mL). Cultures were allowed to grow for 24 hours at 37°C and diluted again to 1:100 in fresh media. This was repeated for 30 days. Each day, each independently evolving population was plated and serially diluted. CFUs were counted (3 plates per sample) and the emergence of non-capsulated mutants was recorded. Noncapsulated mutants are easily visualized by the naked eye as mutants produce smaller, rough and translucent colonies.

### Phage experiments

*(i) Growth curves:* 200 μL of diluted overnight cultures of *Klebsiella spp*. (1:100 in fresh LB) were distributed in a 96-well plate. Cultures were allowed to reach OD = 0.2 and either mitomycin C to 1 μg/mL or 20 μl of PEG-precipitated induced and filtered supernatants at 2 x 10^8^ PFU/mL was added. Growth was then monitored until late stationary phase. *(ii) PEG-precipitation of phages.* Overnight cultures were diluted 1:500 in fresh LB and allowed to grow until OD = 0.2. Mitomycin C was added to final 5 μg/mL. After 4h hours at 37°C, cultures were centrifuged at 4000 rpm and the supernatant was filtered through 0.22um. Filtered supernatants were mixed with chilled PEG-NaCl 5X (PEG 8000 20% and 2.5M of NaCl) and mixed through inversion. Phages were allowed to precipitate for 15 min and pelleted by centrifugation 10 min at 13000 rpm at 4°C. The pellets were dissolved in TBS (Tris Buffer Saline, 50 mM Tris-HCl, pH 7.5, 150 mM NaCl). *(iii) Calculating plaque forming units (PFU)*. Overnight cultures of susceptible or tested strains were diluted 1:100 and allowed to grow until OD = 0.8. 250 μL of bacterial cultures were mixed with 3 mL of top agar (0.7% agar) and poured intro prewarmed LB plates. Plates were allowed to dry before spotting serial dilutions of induced PEG-precipitated phages. Plates were left overnight at room temperature and phage plaques were counted. *(iv) Phage adsorption.* Adsorption of phage particles to the cell surface was performed as previously described ^25^. Briefly, each resistant clone was grown until OD ~0.35. One ml of each culture was transferred to separate wells in a 24-well plate, to which 10 μl of filtered phage lysate (*ca*. 5*10^6^ phage particles) was added. The mix was allowed to sit for 2 minutes at room temperature prior to incubation at 37°C for 15 min with shaking at 140 rpm. Phage adsorption was measured by quantifying the free phage remaining in solution, after centrifugation for 10 minutes at 10000rpm, to get rid of bacterial cells. The supernatant was serially diluted and non-adsorbed phage was quantified by spot titer on a bacterial lawn of strain BJ1. Finally, to quantify how much phage was adsorbed, the non-adsorbed phage was substracted from the initial amount of phage added to the culture.

### Sequencing

*(i) Genomes of phage resistant clones*. Single clones were allowed to grow overday in LB supplemented with 0.7mM EDTA, to limit capsule production. We performed DNA extraction with the guanidium thiocyanate method, with few modifications ^59^. RNAse A treatment (37°C, 30min) was performed before DNA precipitation. Each clone (n=15) was sequenced by Illumina with 150pb paired-end reads, yielding approximately 1 Gb of data per clone. The reads were compared to the reference genome using *breseq* v0.33.2, default parameters. *(ii) wcaJ gene.* PCR of wcaJ was performed using the primers that hybridized 150 base pairs upstream and downstream of the wcaJ gene; K2. wcaJ. 150-5 (5’-GGCGTTCCAGCAAGGGTTATC-3’) and K2.wcaJ. 150-3 (5’-ACGTTCGCGCTTAAATGTG-3’), respectively. To allow full coverage of the gene, PCR products were also sequenced with primer K2.wcaJ.inseq-5 (5’-CTGGGTCTTTACAGAGGAATC-3’). PCR products were sequenced by Sanger and analysed using *APe.*

### Capsule quantification

The bacterial capsule was extracted as described in ^60^. Briefly, 500 μL of an overnight culture was adjusted to OD of 2 and mixed with 100 μL of 1% Zwittergent 3-14 detergent in 100 mM citric acid (pH 2.0) and heated at 56°C for 20 minutes. Afterwards, it was centrifuged for 5 min at 14,000 rpm and 300 μL of the supernatant was transferred to a new tube. Absolute ethanol was added to a final concentration of 80% and the tubes were placed on ice for 20 minutes. After a second wash with ethanol at 70%, the pellet was dried and dissolved in 250 μL of distilled water. The pellet was then incubated for 2 hours at 56°C. Polysaccharides were then quantified by measuring the amount of uronic acid, as described in ^61^. A 1,200 μL volume of 0.0125 M tetraborate in concentrated H2SO4 was added to 200 μL of the sample to be tested. The mixture was vigorously vortexed and heated in a boiling-water bath for 5 min. The mixture was allowed to cool, and 20 μL of 0.15% 3-hydroxydiphenol in 0.5% NaOH was added. The tubes were shaken, and 100 μL were transferred to a microtiter plate for absorbance measurements (520 nm). The uronic acid concentration in each sample was determined from a standard curve of glucuronic acid.

### Citrate quantification

24 hour cultures of BJ1, ST14, their coculture, or blank tubes with citrate, were centrifuged for 10 minutes at 4000 rpm and the supernatant was sterilized with 0.22 μm filter, prior to deproteination by centrifugation in an Amicon tube (10 kDa). Citrate concentration was measured using Citrate assay kit (Sigma-Aldrich MAK333).

### Individual-based simulations of bacteria-phage interactions

Simulations were performed based on the model described in ^42^. Briefly, both bacterial cells and phage particles are independent individuals on an environment represented as a twodimensional grid. The environment is simulated as well-mixed, meaning that positions of bacteria and phage are randomized at each iteration. Bacterial death can be intrinsic (e.g., of old age) or explicit (e.g., lysed by phage). Bacteria can resist phage infection by acquiring a mutation that mimics capsule loss (at varying rates, with a varying fitness cost, see results). Upon phage infection, phage can either follow a lytic cycle or a lysogenic one, according to a stochastic decision defined by the parameters LysogenyAlpha and LysogenyKappa, that takes into consideration the density of nearby phages. When lysogenized, bacteria become insensitive to new phage infections, but the integrated prophage can excise (and thus lead to death of this specific cell) at varying frequencies (see results). The simulations we explored are initiated with 10000 bacterial cells, and 1000 phage particles are added into the environment at the beginning of the simulation. For each condition explored (varying the probability of phage induction, the probability of bacteria acquiring a phage resistance mutation, and the cost of this mutation), we performed 30 replicate simulations, each running for 150 iterations. The values presented in the results correspond to the median of the 30 replicate simulations, for each condition. The set of parameters explored, that are relevant for the questions in this study, are shown in Text S1. Other mechanisms that can be simulated in eVIVALDI (e.g, transduction) were not used in these simulations.

## ACKNOWLEDGEMENTS

The sequencing work was made at the Biomics Platform, C2RT, Institut Pasteur, Paris, France, supported by France Génomique (ANR-10-INBS-09) and IBISA.

## FUNDING

This work was funded by an ANR JCJC (Agence national de recherche) grant [ANR 18 CE12 0001 01 ENCAPSULATION] awarded to O.R. The laboratory is funded by a Laboratoire d’Excellence ‘Integrative Biology of Emerging Infectious Diseases’ (grant ANR-10-LABX-62-IBEID) and the FRM [EQU201903007835]. The funders had no role in study design, data collection and interpretation, or the decision to submit the work for publication.

## COMPETING INTERESTS

Authors declare that we do not have any competing interests in relation to the work described.

## SUPPLEMENTAL MATERIAL

**Figure S1.**
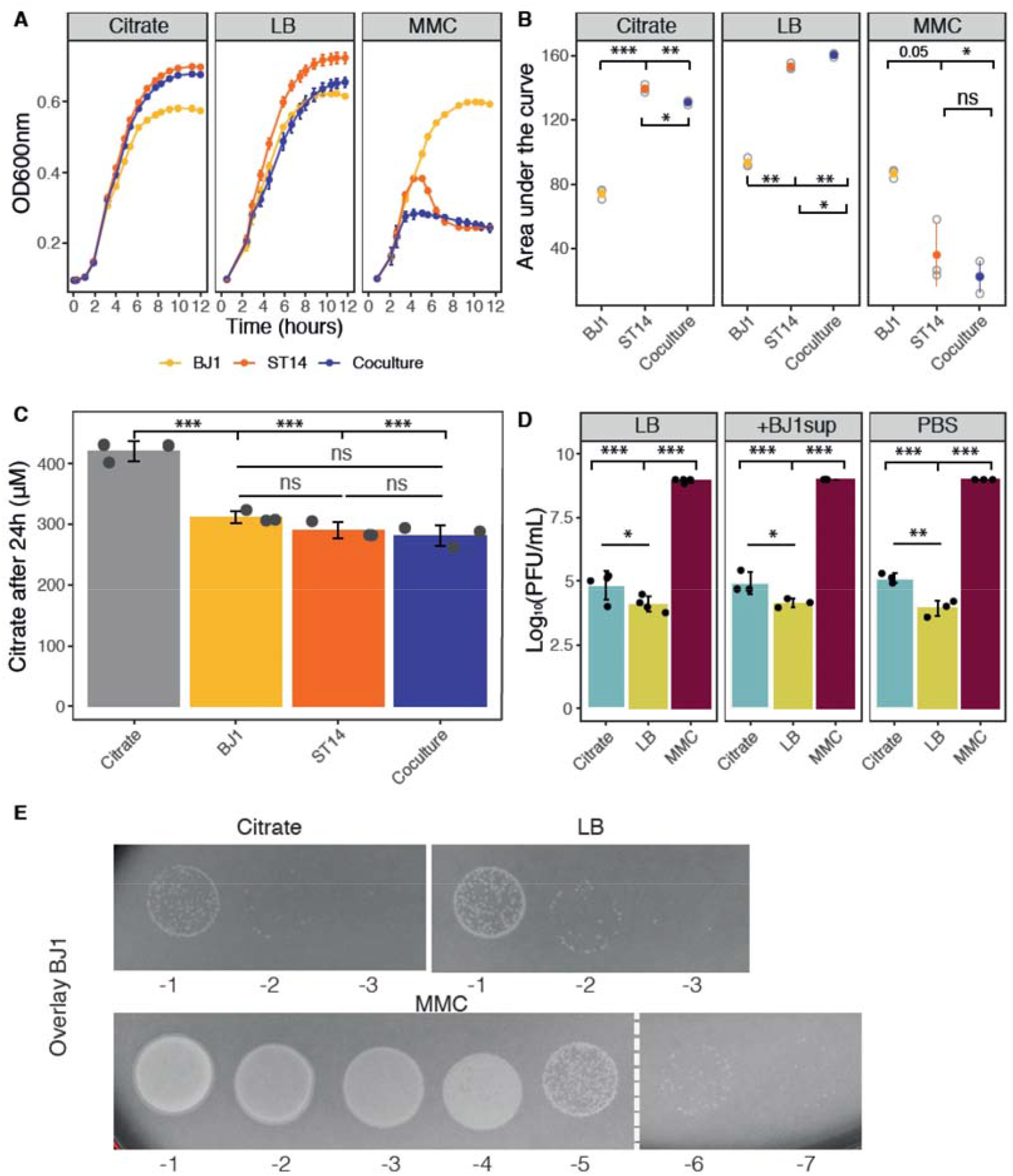
Growth and phage production in experimental conditions. **A.** Growth of each strain and their coculture in each experimental condition. Three independent clones and cocultures were measured. B. Area under the growth curves, calculated with *trapz* function from pracma package for R. **C.** Citrate amount in 24-hour cultures of strain BJ1, ST14, 1:1 coculture and control cultures of plain LB with citrate. Each dot represents an independent experiment. Error bars indicate standard deviation. **D.** Strain ST14 was grown in LB, in LB diluted (1:1) in spent supernatant from BJ1, or in LB diluted (1:1) in PBS. Phage lysates were prepared and PFU analysed on a lawn of strain BJ1. Each dot represents and independent experiment. Error bars indicate standard deviation. **E.** Lysis plaques on an overlay of strain BJ1 after addition of 10 ul of supernatant of strain ST14 grown in the different experimental conditions. * P< 0.05,** P< 0.01,***P<0.001 for ANOVA with Tukey *post hoc* corrections.

**Figure S2.**
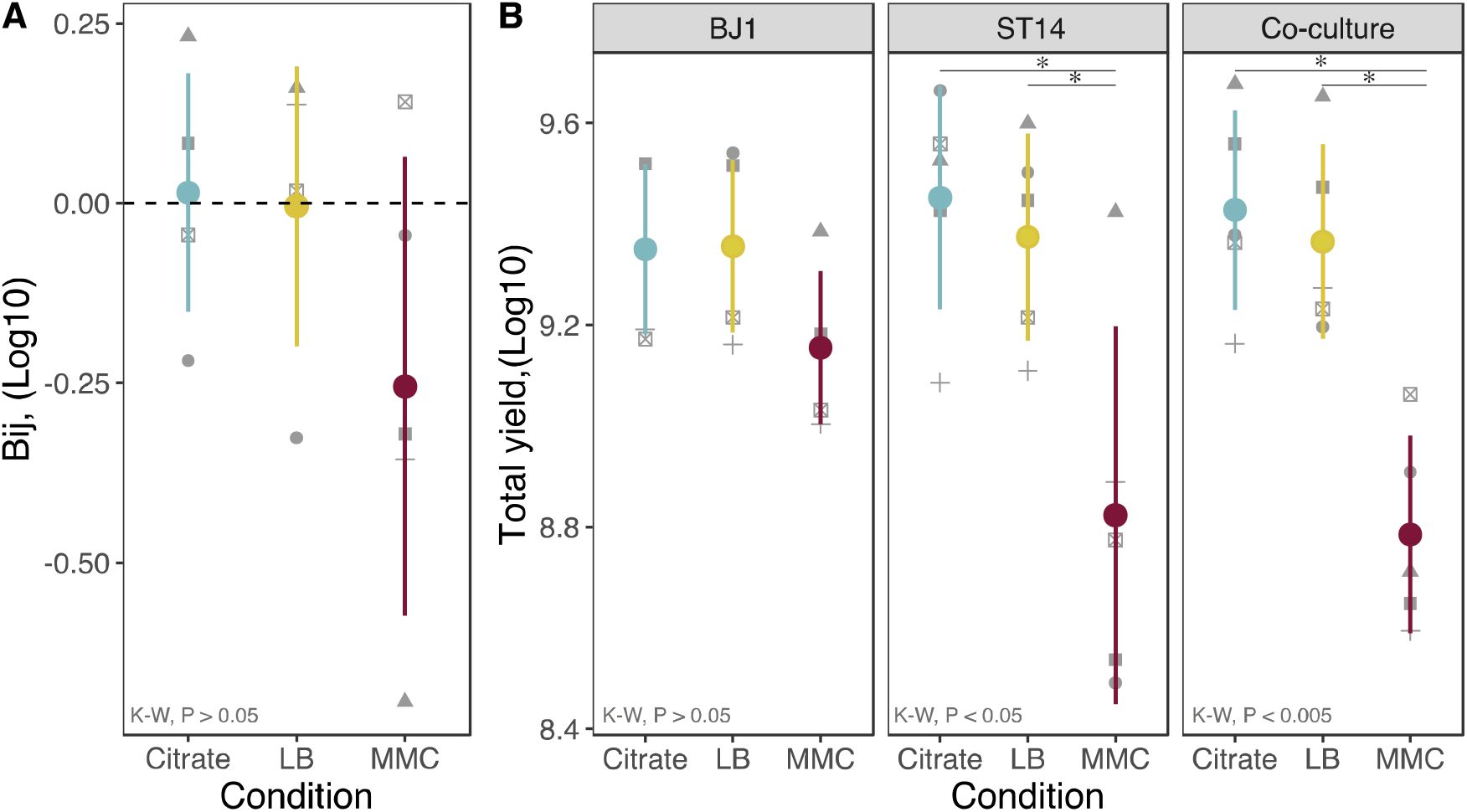
Total growth in the different media. **A.** The effect of mixing on total growth or population yield is expressed as *Bi(j)*, in which positive values represent more growth during coculture than expected from the pure cultures. Each shape represents an independent experiment, N=5. Error bars indicate standard deviation. **B.** The total yield of each culture is expressed as the log10-transformed of CFU/mL. Each shape represents an independent experiment, N=5. P-values correspond to Kruskal-Wallis(K-W) rank sum test, followed by pairwise Wilcoxon test for differences across conditions, with Benjamini-Hochberg correction. * P< 0.05

**Figure S3.**
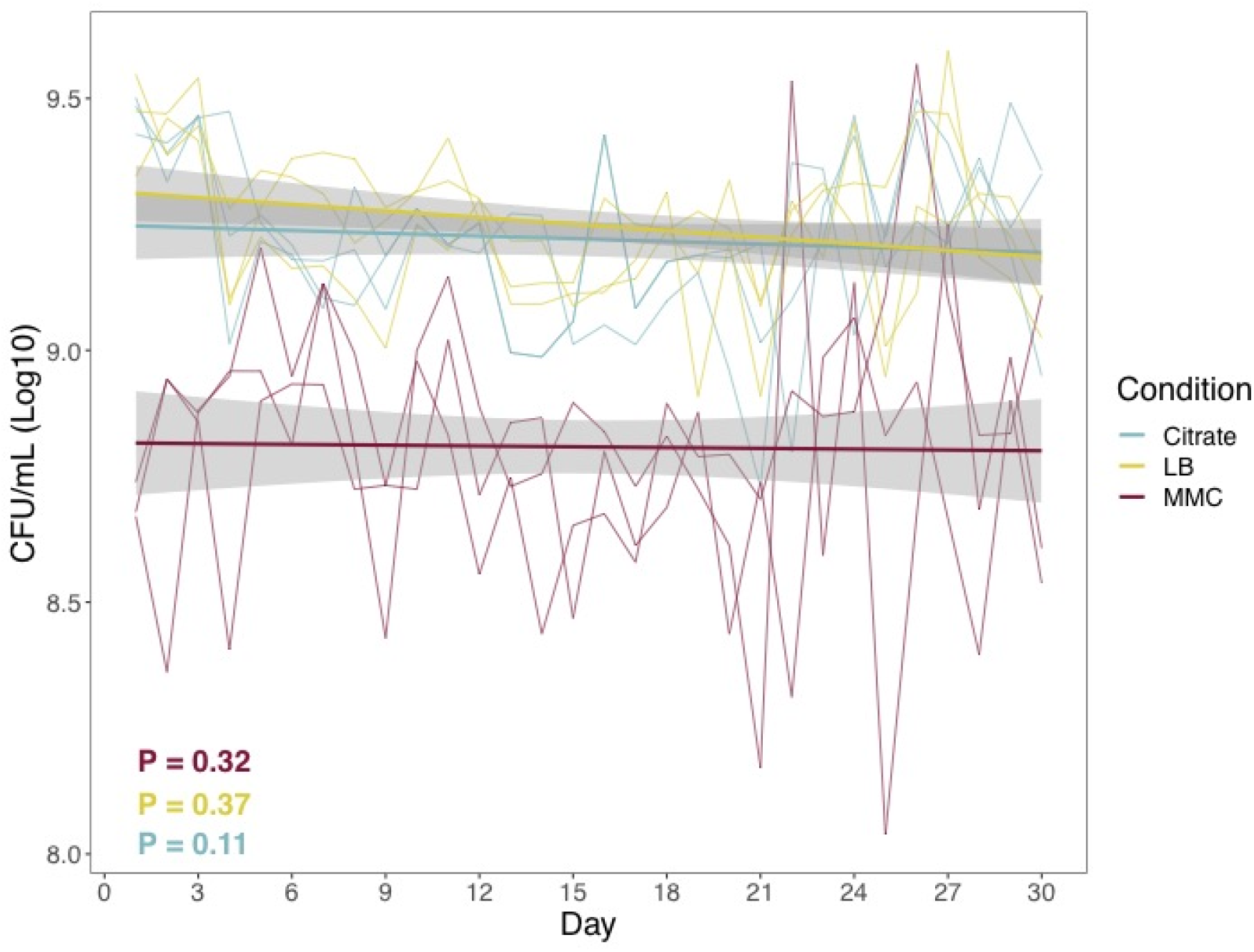
Total number of cells across evolving populations. Each independent line represents an independently evolving population. Bold lines represent regression line from a linear model. The slopes of regression model were calculated for each independent population, and tested with a one-sample t-test for significant difference from 0.

**Figure S4.**
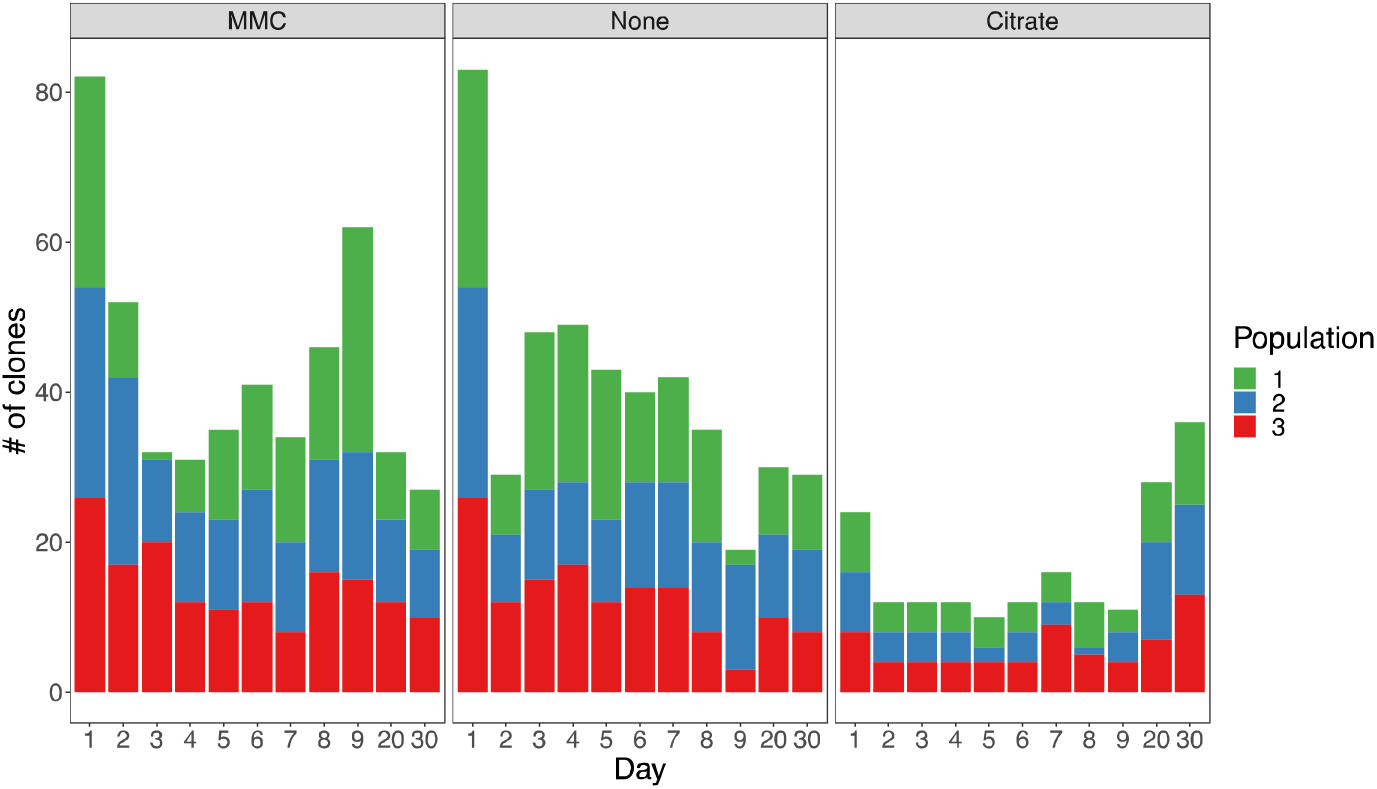
Number of clones from strain BJ1 analysed each day from each population. Each day a total of 93 capsulated clones from every population in each evolutionary treatment were isolated, except for day 1, when 186 capsulated clones (93 x 2) were isolated. The number of 93 was chosen for it to conveniently fit in a 96-well plate and allowing three wells for internal controls (ancestors and blank).

**Figure S5.**
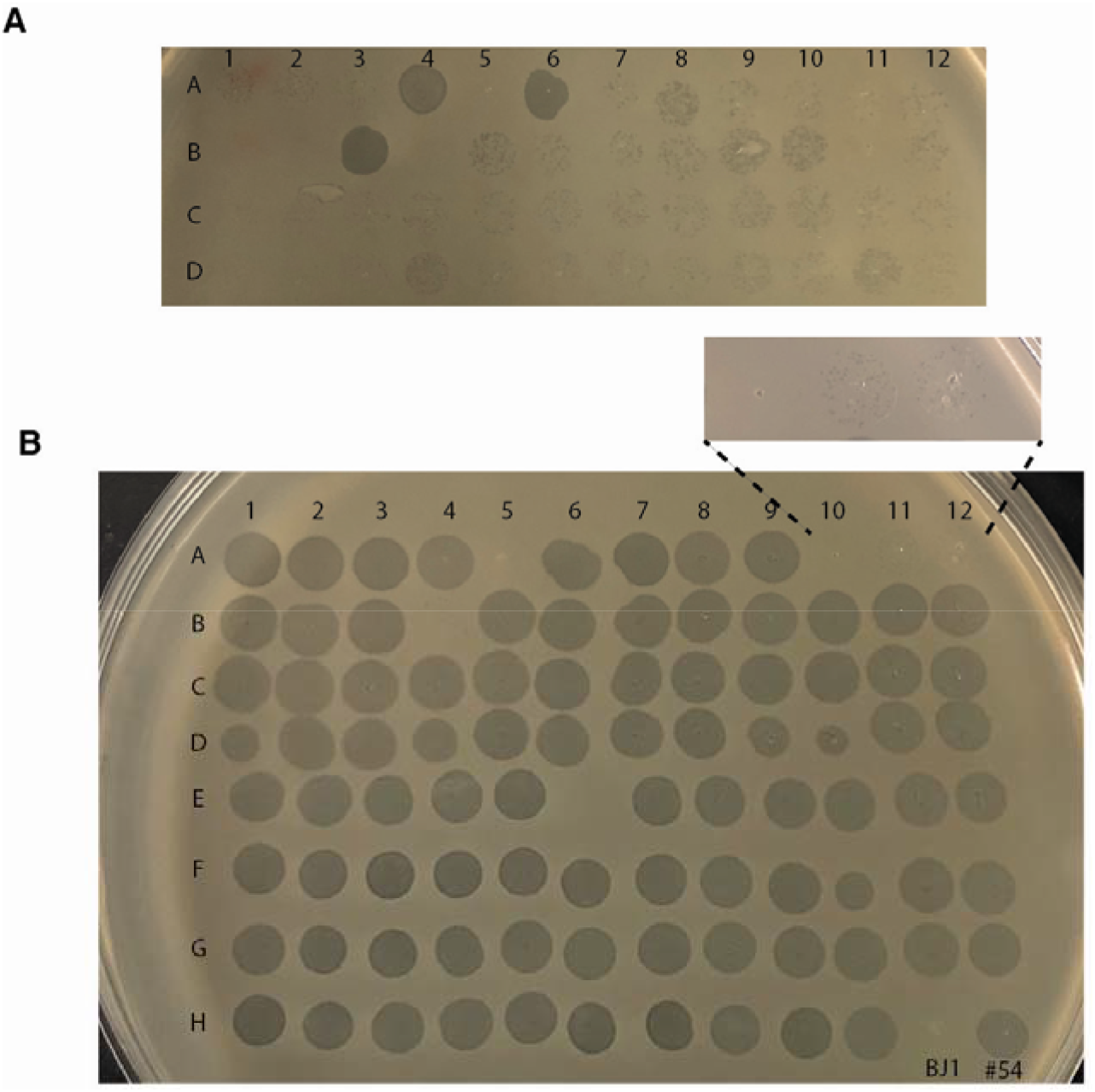
Lysogens can release viable phages that infect the ancestral BJ1 strain. 93 lysogens were grown overnight in 100 μL of LB in a 96-well microtiter plate. Cultures were then diluted 1:100, and allowed to grow for 6 hours (**A**, non induced lysogens), or two hours, after which, MMC was added to a final concentration of 5 μg/mL (**B**, induced lysogens). Cultures were allowed to grow for 4 more hours. After a total of 6 hours of overday growth, the microtiter plate was centrifuged to allow cells to pellet, and 4 μL of the supernatant was spotted on an overlay of 0.7% Lennox agar with the ancestral BJ1 strain. Two controls are included; ancestral BJ1(well H11) and ancestral ST14 (well H12).

**Figure S6.**
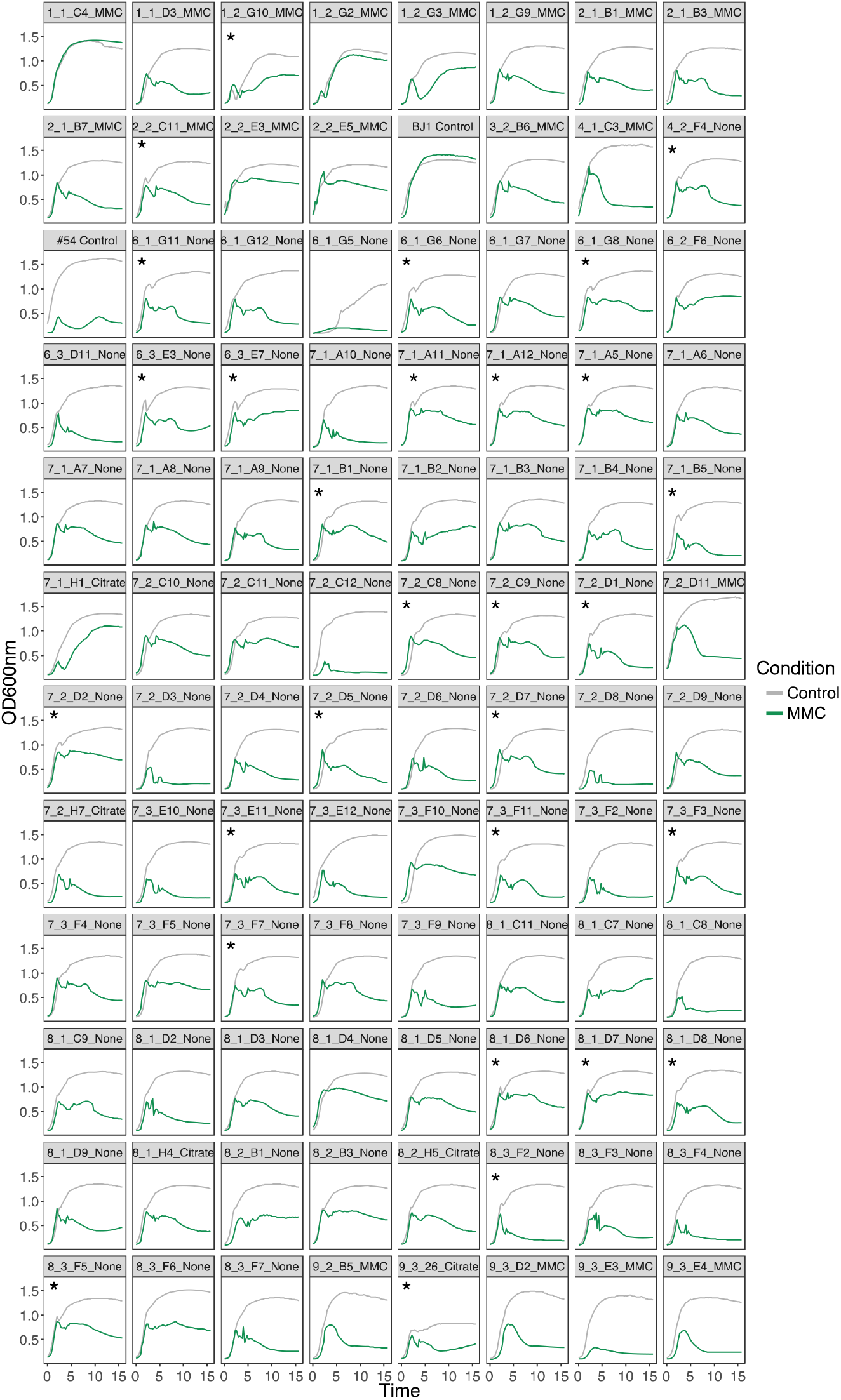
Growth of newly lysogenized clones. Clones were grown in either LB (control, grey line) for 16 hours or in LB to which MMC (1 μg/ml) was added after 1.5 hours (green). Labels on top of each graph correspond to the isolation day, population number, clone ID and condition in which it evolved. Star (*) indicates clones in which death is observed at the end of exponential phase, suggestive of unusually large phage outburst. Only one replicate per curve is shown for clarity purposes.

**Figure S7.**
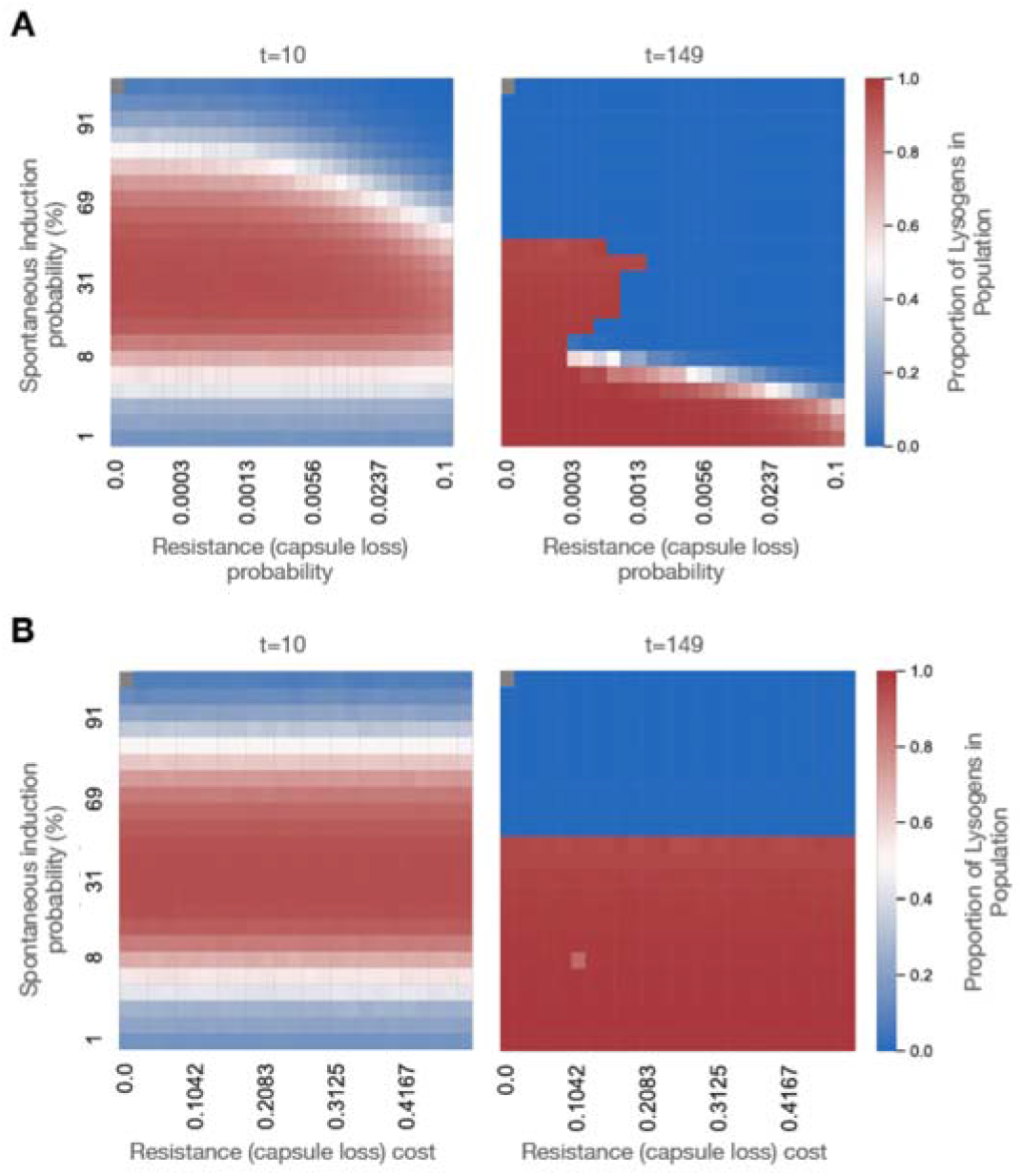
Simulated temporal dynamics of the competition between different phage resistance mechanisms. eVIVALDI was used to simulate a scenario where an initial population of phage-sensitive bacteria (n=#) is co-inoculated with # temperate phages. Upon infection, bacteria either die from the phage’s lytic cycle or become lysogens if the phage follows the lysogenic cycle. Integrated phage (prophages) can subsequently spontaneously induce and restart the lytic cycle, leading to bacterial death. Bacteria can also become phage-resistant (before or after becoming lysogens) by random mutation that represents the loss of the capsule. **A.** Heatmap of the proportion of lysogens in the populations at intermediate (t=10) and final (t=149) timepoints, as a function of the spontaneous induction probability (y-axis) and the probability of phage resistant (capsule-less) mutants to emerge. The fitness cost of the resistant mutants is fixed at 10%. **B.** Heatmap of the proportion of lysogens in the populations at intermediate (t=10) and final (t=149) timepoints, as a function of the spontaneous induction probability (y-axis) and the fitness cost of a phage resistant mutation (loss of capsule). The probability of these mutations to emerge is fixed at 0.001. For B, C and D, dark red colors indicate a prevalence of lysogens in the population, while deep blue colors indicate their absence. Each square in the heatmaps represents the median of the 20 independent replicate simulations for each combination of parameters.

**Figure S8.**
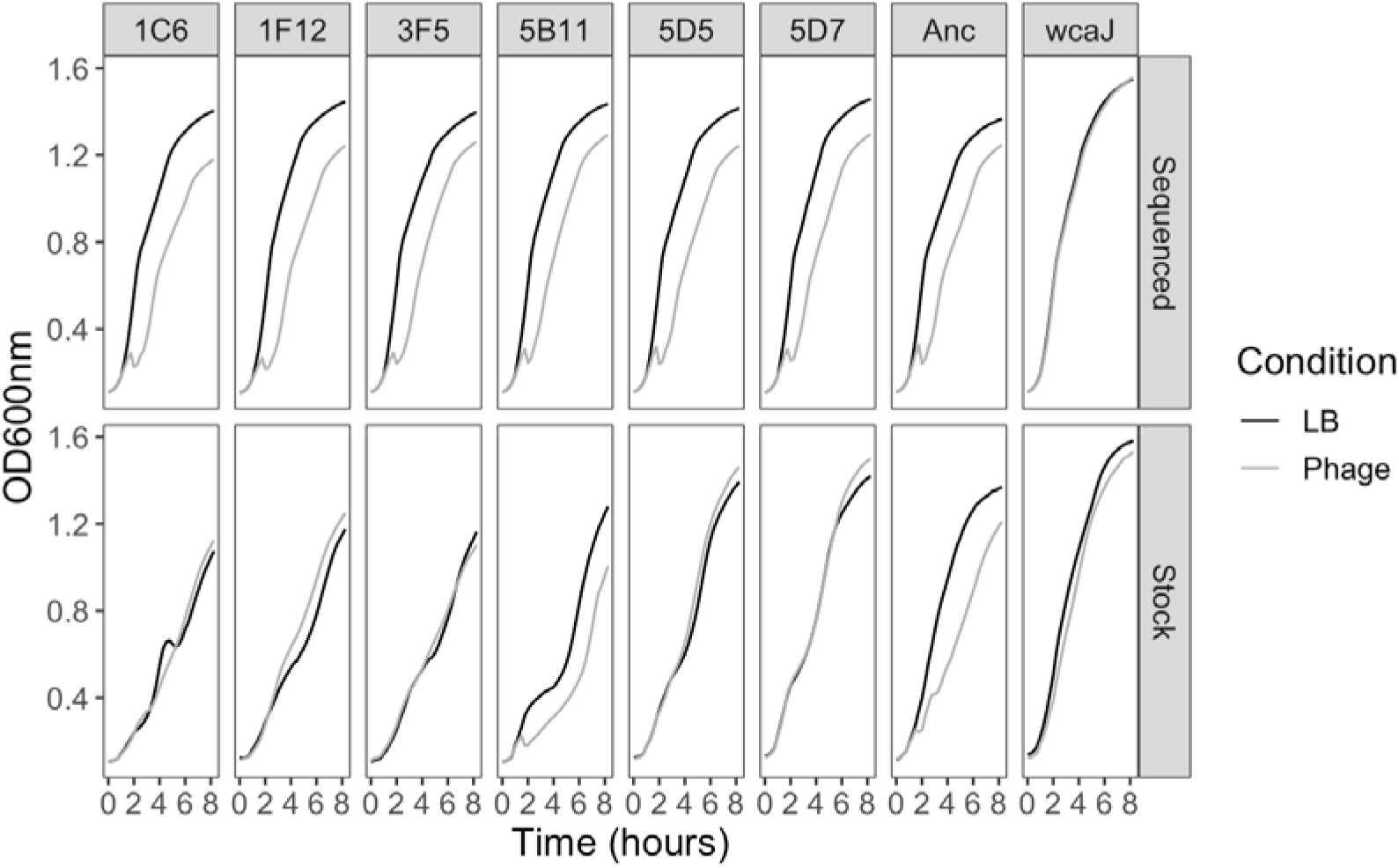
Growth curves of resistant clones with no fixed mutations. Growth of resistant clones from the original stock were directly initiated from the glycerol stock without performing a preconditioning culture. Sequenced clones underwent two extra rounds of culture since the original stock; restreaked on agar and an overnight LB culture used to extract the genome and to generate a new glycerol stock, from which all other experiments were initiated. N=4. No error bars are shown for clarity purposes.

**Figure S9.**
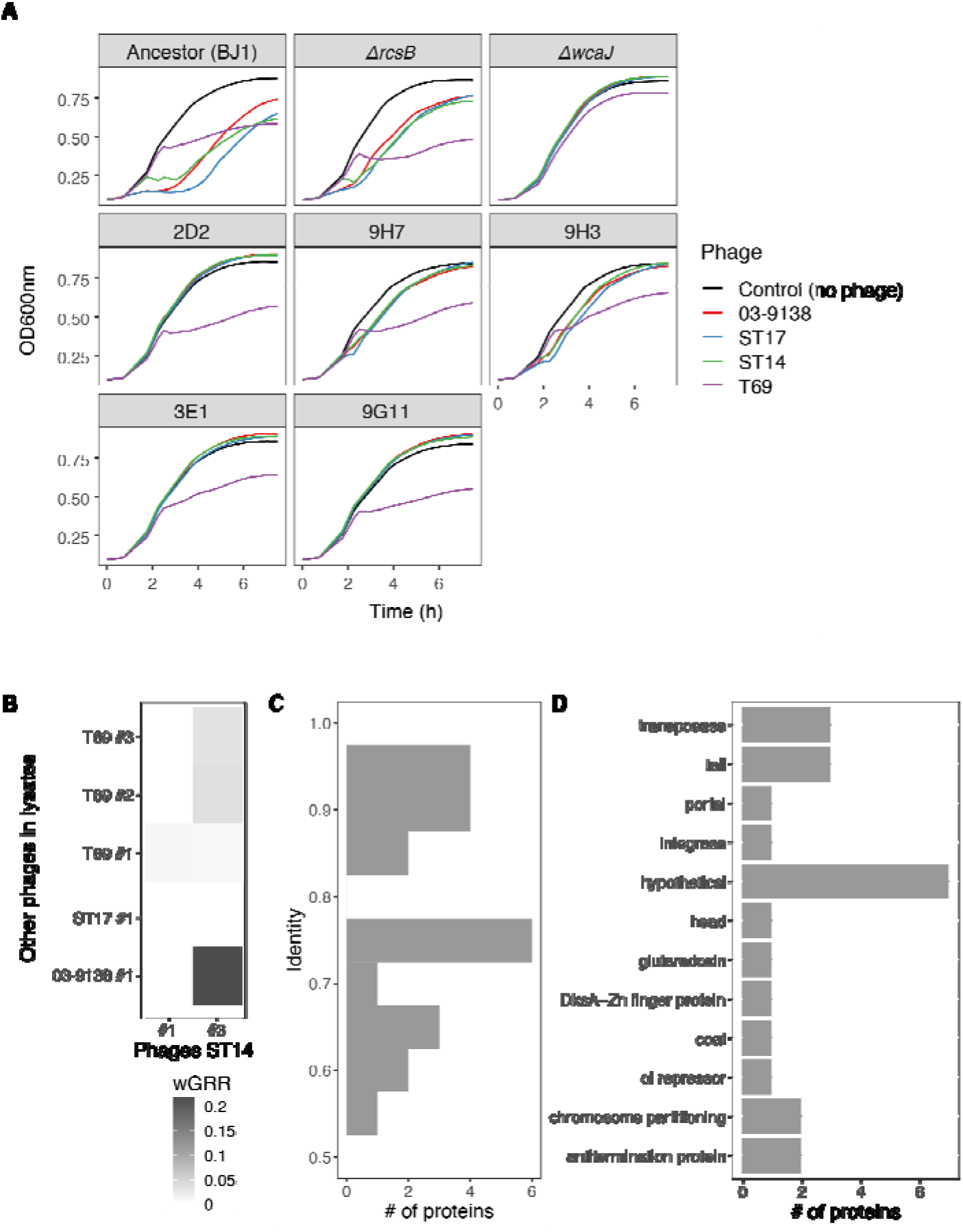
Characterization of lysates produced by other K2 strains. **A**. Growth curves of resistant mutants and control strains in the presence of lysates produced by other K2 strains. Error bars are not shown to improve visibility. **B**. Phage simimarity as expressed by wGRR of intact phages in lysates. **C**. Identity of proteins in intact phages present in other lysates against proteins in phages of strain ST14. (Only values > 0.5 as determined by blastp are displayed). **D.** Predicted function of proteins with an identity > 0.5. The functional characterization was performed by annotating the intact phages with prokka v1.14.0 ^62^, and pVOG profiles ^63^ searched for using HMMER v3.2 ^64^. For each protein, the pVOG with the lowest p-value was kept, and keywords analyzed.

## SUPPLEMENTAL TABLES

**Table S1.** PHASTER prophage prediction in strain ST14. The genome was analysed with PHASTER ^65^ in April 2021.

| Phage # | Length (kb) | Classification | Coding proteins | Comment |
| --- | --- | --- | --- | --- |
| 1 | 108.9 | intact | integrase, recombinase, terminase, capsid, tail, lysin | Can lysogenize BJ1<sub>24</sub> |
| 2 | 17.2 | incomplete | integrase, head, tail | P2 remnant? |
| 3 | 52.8 | intact | tail, head, terminase, transposase, lysis, integrase | Can lysogenize BJ1<sub>24</sub> |
| 4 | 45.5 | intact | tail, capsid, terminase, head |  |
| 5 | 46.4 | intact | integrase, tail, terminase, head, protease, capsid |  |
| 6 | 47.6 | questionable | integrase, recombinase, tail, terminase, head, coat |  |
| 7 | 12.9 | incomplete | tail genes | Most likely not functional. No packaging genes |
| 8 | 12.7 | incomplete | tail genes | Most likely not functional. No packaging genes |
| 9 | 8.7 | incomplete | recombinase, transposase | Most likely not functional. |
| 10 | 10.2 | incomplete | transposase, tail, portal, terminase | Most likely not functional |

**Table S2.**
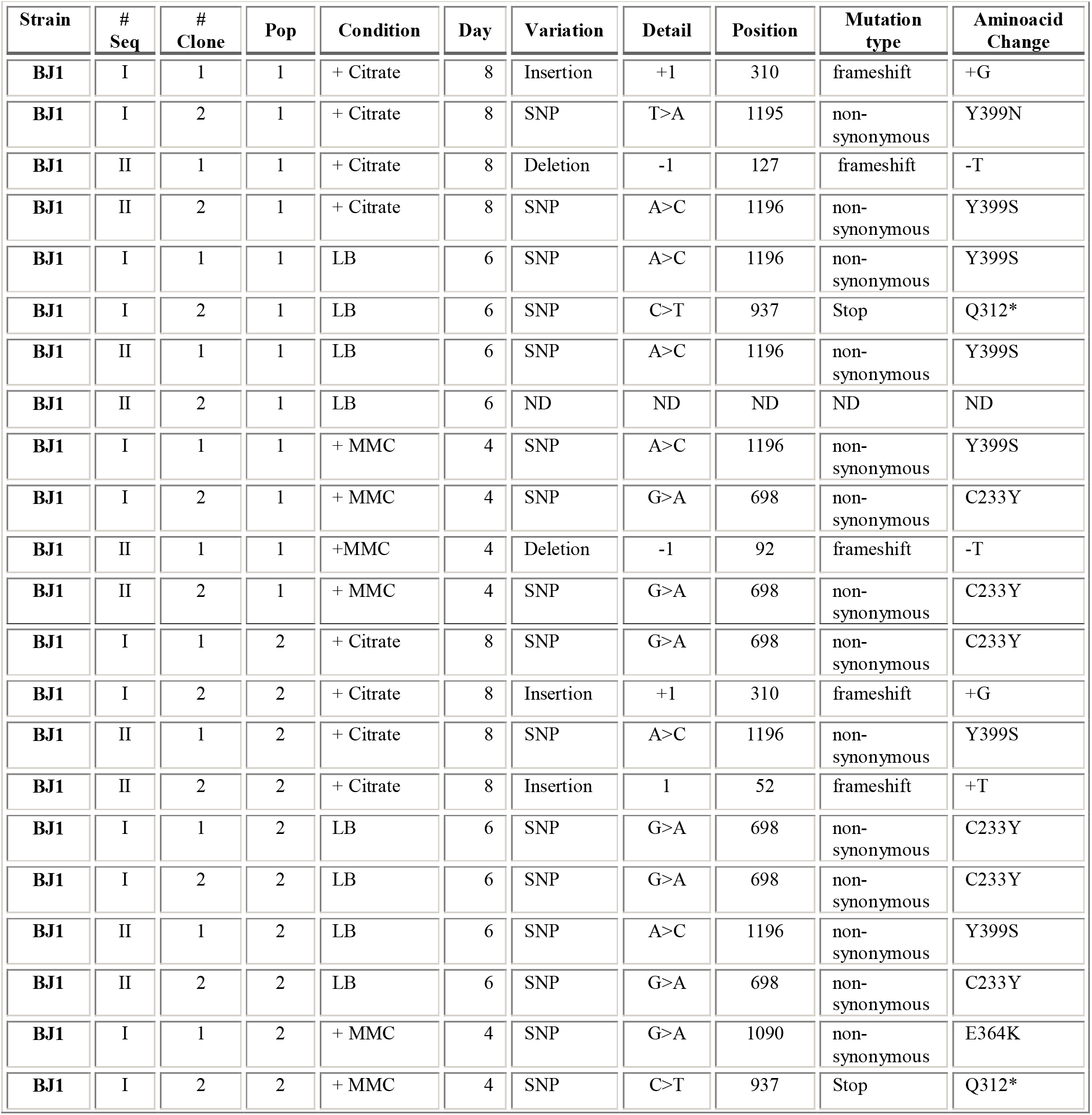
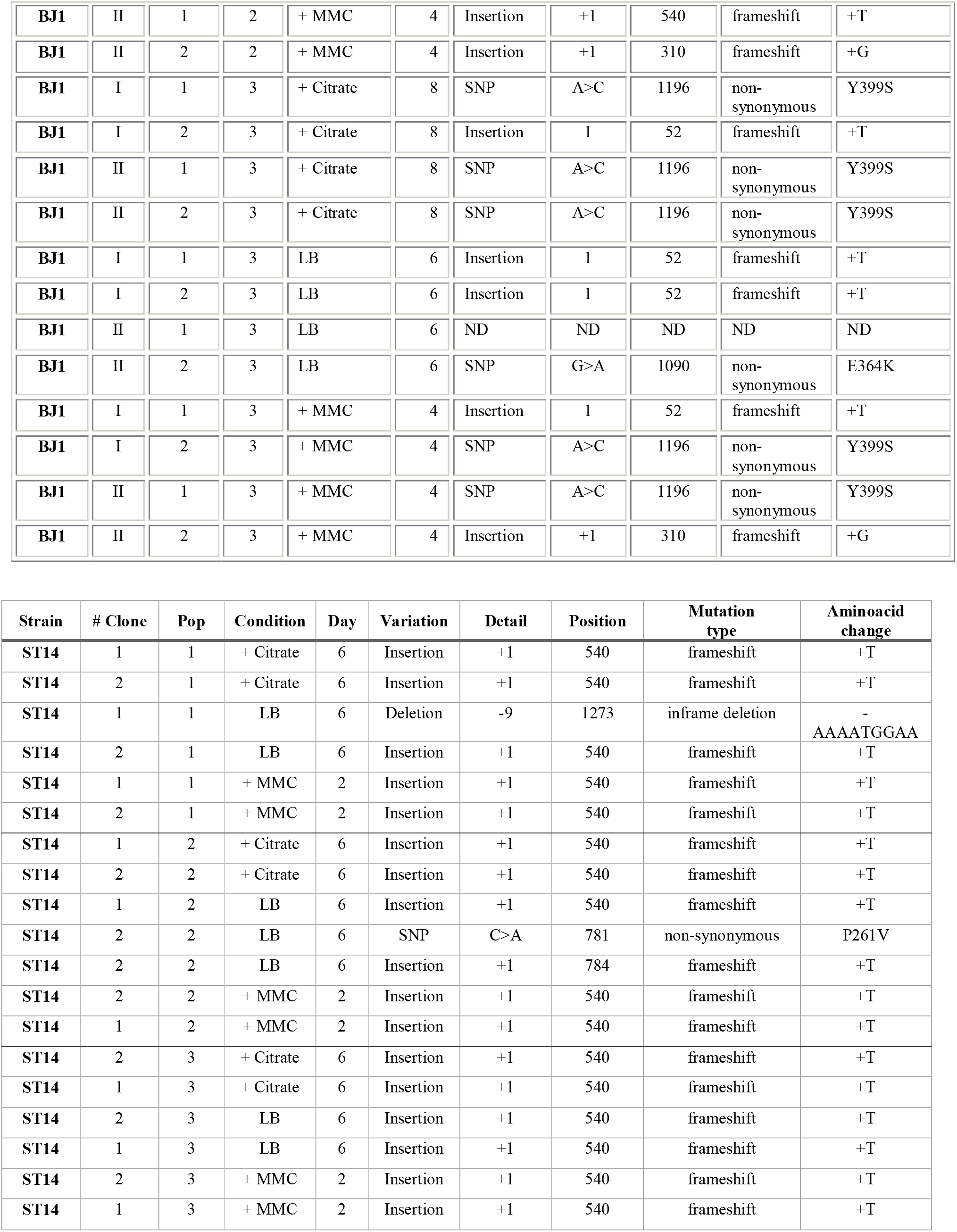
Mutations accumulated in *wcaJ* gene resulting in non-capsulated mutants. ND; none detected. At the day in which at least 50% of the clones of each independently evolving population were non-capsulated, two of such clones were isolated (# Clone), the *wcaJ* gene amplified by PCR and sequenced by Sanger. For BJ1, this was repeated twice independently (# Seq) (N=36 for BJ1 and N=18 for ST14). Independently evolving populations (Pop) are identified with number from 1 to 3. Sequencing of ancestor revealed that no mutations were present in the *wcaJ* genes.

| Strain | # Seq | # Clone | Pop | Condition | Day | Variation | Detail | Position | Mutation type | Aminoacid Change |
| --- | --- | --- | --- | --- | --- | --- | --- | --- | --- | --- |
| BJ1 | I | 1 | 1 | + Citrate | 8 | Insertion | +1 | 310 | frameshift | +G |
| BJ1 | I | 2 | 1 | + Citrate | 8 | SNP | T>A | 1195 | non-synonymous | Y399N |
| BJ1 | II | 1 | 1 | + Citrate | 8 | Deletion | -1 | 127 | frameshift | -T |
| BJ1 | II | 2 | 1 | + Citrate | 8 | SNP | A>C | 1196 | non-synonymous | Y399S |
| BJ1 | I | 1 | 1 | LB | 6 | SNP | A>C | 1196 | non-synonymous | Y399S |
| BJ1 | I | 2 | 1 | LB | 6 | SNP | C>T | 937 | Stop | Q312* |
| BJ1 | II | 1 | 1 | LB | 6 | SNP | A>C | 1196 | non-synonymous | Y399S |
| BJ1 | II | 2 | 1 | LB | 6 | ND | ND | ND | ND | ND |
| BJ1 | I | 1 | 1 | + MMC | 4 | SNP | A>C | 1196 | non-synonymous | Y399S |
| BJ1 | I | 2 | 1 | + MMC | 4 | SNP | G>A | 698 | non-synonymous | C233Y |
| BJ1 | II | 1 | 1 | +MMC | 4 | Deletion | -1 | 92 | frameshift | -T |
| BJ1 | II | 2 | 1 | + MMC | 4 | SNP | G>A | 698 | non-synonymous | C233Y |
| BJ1 | I | 1 | 2 | + Citrate | 8 | SNP | G>A | 698 | non-synonymous | C233Y |
| BJ1 | I | 2 | 2 | + Citrate | 8 | Insertion | +1 | 310 | frameshift | +G |
| BJ1 | II | 1 | 2 | + Citrate | 8 | SNP | A>C | 1196 | non-synonymous | Y399S |
| BJ1 | II | 2 | 2 | + Citrate | 8 | Insertion | 1 | 52 | frameshift | +T |
| BJ1 | I | 1 | 2 | LB | 6 | SNP | G>A | 698 | non-synonymous | C233Y |
| BJ1 | I | 2 | 2 | LB | 6 | SNP | G>A | 698 | non-synonymous | C233Y |
| BJ1 | II | 1 | 2 | LB | 6 | SNP | A>C | 1196 | non-synonymous | Y399S |
| BJ1 | II | 2 | 2 | LB | 6 | SNP | G>A | 698 | non-synonymous | C233Y |
| BJ1 | I | 1 | 2 | + MMC | 4 | SNP | G>A | 1090 | non-synonymous | E364K |
| BJ1 | I | 2 | 2 | + MMC | 4 | SNP | C>T | 937 | Stop | Q312* |
| **BJ1** | II | 1 | 2 | + MMC | 4 | Insertion | +1 | 540 | frameshift | +T |
| **BJ1** | II | 2 | 2 | + MMC | 4 | Insertion | +1 | 310 | frameshift | +G |
| **BJ1** | I | 1 | 3 | + Citrate | 8 | SNP | A>C | 1196 | non-synonymous | Y399S |
| **BJ1** | I | 2 | 3 | + Citrate | 8 | Insertion | 1 | 52 | frameshift | +T |
| **BJ1** | II | 1 | 3 | + Citrate | 8 | SNP | A>C | 1196 | non-synonymous | Y399S |
| **BJ1** | II | 2 | 3 | + Citrate | 8 | SNP | A>C | 1196 | non-synonymous | Y399S |
| **BJ1** | I | 1 | 3 | LB | 6 | Insertion | 1 | 52 | frameshift | +T |
| **BJ1** | I | 2 | 3 | LB | 6 | Insertion | 1 | 52 | frameshift | +T |
| **BJ1** | II | 1 | 3 | LB | 6 | ND | ND | ND | ND | ND |
| **BJ1** | II | 2 | 3 | LB | 6 | SNP | G>A | 1090 | non-synonymous | E364K |
| **BJ1** | I | 1 | 3 | + MMC | 4 | Insertion | 1 | 52 | frameshift | +T |
| **BJ1** | I | 2 | 3 | + MMC | 4 | SNP | A>C | 1196 | non-synonymous | Y399S |
| **BJ1** | II | 1 | 3 | + MMC | 4 | SNP | A>C | 1196 | non-synonymous | Y399S |
| **BJ1** | II | 2 | 3 | + MMC | 4 | Insertion | +1 | 310 | frameshift | +G |

| **Strain** | **# Clone** | **Pop** | **Condition** | **Day** | **Variation** | **Detail** | **Position** | **Mutation type** | **Aminoacid change** |
| --- | --- | --- | --- | --- | --- | --- | --- | --- | --- |
| **ST14** | 1 | 1 | + Citrate | 6 | Insertion | +1 | 540 | frameshift | +T |
| **ST14** | 2 | 1 | + Citrate | 6 | Insertion | +1 | 540 | frameshift | +T |
| **ST14** | 1 | 1 | LB | 6 | Deletion | -9 | 1273 | inframe deletion | -<br>AAAATGGAA |
| **ST14** | 2 | 1 | LB | 6 | Insertion | +1 | 540 | frameshift | +T |
| **ST14** | 1 | 1 | + MMC | 2 | Insertion | +1 | 540 | frameshift | +T |
| **ST14** | 2 | 1 | + MMC | 2 | Insertion | +1 | 540 | frameshift | +T |
| **ST14** | 1 | 2 | + Citrate | 6 | Insertion | +1 | 540 | frameshift | +T |
| **ST14** | 2 | 2 | + Citrate | 6 | Insertion | +1 | 540 | frameshift | +T |
| **ST14** | 1 | 2 | LB | 6 | Insertion | +1 | 540 | frameshift | +T |
| **ST14** | 2 | 2 | LB | 6 | SNP | C>A | 781 | non-synonymous | P261V |
| **ST14** | 2 | 2 | LB | 6 | Insertion | +1 | 784 | frameshift | +T |
| **ST14** | 2 | 2 | + MMC | 2 | Insertion | +1 | 540 | frameshift | +T |
| **ST14** | 1 | 2 | + MMC | 2 | Insertion | +1 | 540 | frameshift | +T |
| **ST14** | 2 | 3 | + Citrate | 6 | Insertion | +1 | 540 | frameshift | +T |
| **ST14** | 1 | 3 | + Citrate | 6 | Insertion | +1 | 540 | frameshift | +T |
| **ST14** | 2 | 3 | LB | 6 | Insertion | +1 | 540 | frameshift | +T |
| **ST14** | 1 | 3 | LB | 6 | Insertion | +1 | 540 | frameshift | +T |
| **ST14** | 2 | 3 | + MMC | 2 | Insertion | +1 | 540 | frameshift | +T |
| **ST14** | 1 | 3 | + MMC | 2 | Insertion | +1 | 540 | frameshift | +T |

## Notes

### Competing Interest Statement

The authors have declared no competing interest.

### Summary of Updates

Figure 3: added a new panel (panel C) Figure 6: added panel B. Introduction and discussion revised

